# TTL proteins scaffold brassinosteroid signaling components at the plasma membrane to optimize signal transduction in plant cells

**DOI:** 10.1101/351056

**Authors:** Vítor Amorim-Silva, Álvaro García-Moreno, Araceli G. Castillo, Naoufal Lakhssassi, Jessica Pérez-Sancho, Yansha Li, Alicia Esteban del Valle, David Posé, Josefa Pérez-Rodriguez, Jinxing Lin, Victoriano Valpuesta, Omar Borsani, Cyril Zipfel, Alberto P. Macho, Miguel A. Botella

## Abstract

Brassinosteroids (BRs) form a group of steroidal hormones essential for plant growth, development and stress responses. Here, we report that plant-specific TETRATRICOPEPTIDE THIOREDOXIN-LIKE (TTL) proteins are positive regulators of BR signaling functioning as scaffold for BR signaling components in Arabidopsis. TTL3 forms a complex with all core components involved in BR signaling, including the receptor kinase BRASSINOSTEROID INSENSITIVE1 (BRI1), the transcription factor BRASSINAZOLE RESISTANT1 (BZR1) and the phosphatase BRI1-SUPPRESSOR1 (BSU1), but excluding the co-receptor BAK1. TTL3 is mainly localized in the cytoplasm, but BR treatment increases its localization at the plasma membrane, where it strengthens the association with BR signaling components. Consistent with a role in BR signaling, mutations in *TTL3* and related *TTL1* and *TTL4* genes cause reduced BR responsiveness. We propose a mechanistic model for BR signaling, in which cytoplasmic/nuclear BR components bound to TTL proteins are recruited to the plasma membrane upon BR perception, which in turn allows the assembly of a BR signaling complex, leading to the de-phosphorylation and nuclear accumulation of the transcription factors BZR1 and BES1.

## Introduction

Plants live in constantly changing environments that are often unfavorable or stressful for growth and development. In these conditions it is essential to balance growth and stress responses to ensure proper allocation of resources ^1^. While an active growth causes the generation of new roots and leaves, allowing a better exploitation of environmental resources, it can also cause the depletion of resources that could be important for the survival under stress episodes ^23^. Brassinosteroids (BRs) are a family of growth-promoting hormones having essential roles in a wide range of developmental and physiological processes ^2,4,5^ However, in addition to their well-established function in growth, essential roles in the trade-off between growth and tolerance to biotic and abiotic stress episodes are now being unveiled ^6–9^

BRs are perceived at the plasma membrane by ligand-induced heterodimers of the receptors kinases BRASSINOSTEROID INSENSITIVE1 (BRI1) and SOMATIC EMBBRYOGENESIS RECEPTOR KINASE (SERK) protein family-members, which activates an interconnected signal transduction cascade, leading to the transcriptional regulation of BR-responsive genes ^5^. BRI1 KINASE INHIBITOR 1 (BKI1) dissociates from activated BRI1, which phosphorylates the kinases BR-SIGNALING KINASE1 (BSK1) and the CONSTITUTIVE DIFFERENTIAL GROWTH1 (CDG1), which in turn phosphorylate the phosphatase BRI1-SUPPRESSOR1 (BSU1). Then, the active (phosphorylated) BSU1 lead to dephosphorylation and inactivation of the glycogen synthase kinase 3 (GSK3)-like BRASSINOSTEROID INSENSITIVE2 (BIN2), a key regulator in BR signaling. In the absence of BRs, BIN2 is active and phosphorylates the two homologous transcription factors BRASSINAZOLE RESISTANT1 (BZR1) and BRI1-ETHYL METHANESULFONATE SUPPRESSOR 1 (BES1/BZR2), which results in their inactivation and degradation. In contrast, when BR is present, BIN2 is inactivated and degraded by the proteasome, which leads to both the stabilization and activation of BZR1 and BES1, and therefore to transcriptional regulation of BR-responsive genes ^5,10^.

In Arabidopsis, the *TETRATRICOPEPTIDE THIOREDOXIN-LIKE (TTL)* gene family is composed of four members (*TTL1* to *TTL4*) and mutations in *TTL1, TTL3*, and *TTL4* genes cause reduced growth under abiotic stresses such as salinity and drought ^11–13^. This stress hypersensitivity is exacerbated in double and triple *ttl* mutants ^12^ The *TTL2* gene is specifically expressed in pollen grains and does not have a role in stress tolerance, but it is important for male sporogenesis ^12^ *TTL* genes encode proteins with a common modular architecture containing six Tetratricopeptide Repeat (TPR) domains distributed in specific positions throughout the sequence and a C-terminal sequence with homology to thioredoxins ^11,12^ TPR domains are well-described protein-protein interaction modules, however how TTL proteins function mechanistically in stress tolerance remains elusive.

Several evidences point to a role of TTL proteins in BR responses, which open the possibility of a direct link between stress tolerance and BR-signaling by the TTL proteins. First, the TTL3 protein, whose gene is the most expressed among the *TTL* gene family, was identified as an interacting partner of the activated (phosphorylated) cytoplasmic domain of VASCULAR HIGHWAY1/BRI1-LIKE RECEPTOR KINASE2 (BRL2). Although BRL2 cannot bind BRs (Belkhadir, 2015), it is a receptor-like kinase homologous to BRI1 with a role in vascular development ^13^ Second, a *ttl3* mutant showed altered growth in the presence of exogenous BRs ^13^. Third, TTL proteins are predicted to interact and function as co-chaperones of Hsp90 ^14^, which has been recently identified to have important roles in BR signaling by interacting with specific BR signaling components ^15–18^. Fourth, a triple *ttl1 ttl3 ttl4* mutant in *TTL1, TTL3*, and *TTL4* shows defects in vasculature development and male sporogenesis, hallmarks of BR defective mutants ^12,19^. Finally, *TTL1, TTL3*, and *TTL4* genes are specifically induced by BR application but not by other hormones ^14^.

Based on phenotypic and molecular analyses we show that *TTL1, TTL3*, and *TTL4* genes, in addition to their reported role in abiotic stress tolerance, are positive regulators of BR signaling. The well-described TPR protein interaction modules of TTL proteins and their role in the assembly of multiprotein complexes ^20–22^ led us to hypothesize that these proteins could function as scaffold for BR signaling. Indeed, we show that TTL3 interacts with BRI1, BSU1 and BZR1 and associates *in vivo* with the majority of BR signaling components but not with BAK1. We also show that a functional TTL3 tagged with a Green Fluorescent Protein (GFP) shows a dual cytoplasmic and plasma membrane localization that is dependent on endogenous BR content. Furthermore TTL3 highly enhances the interaction between BSK1 and BZR1. Taking together these results, we reveal that TTL proteins function in BR-regulated stress tolerance in plants and propose a model in which TTL proteins function in optimizing BR signal transduction by acting as a scaffold of BR signaling components.

## Results

### TTL3 interacts with a BAKI-independent phosphomimetic BRI1 mutant

The TTL3 protein (also known as VIT1) has been identified as an interactor of the activated (phosphorylated) cytoplasmic domain of BRL2 ^13^, a receptor kinase of the BRI1 family with a role in vascular development ^13,23^ *TTL3* belongs to a family of 4 genes (from *TTL1* to *TTL4*) in *Arabidopsis* ^11,12^. We confirmed defects in vein formation using a different *ttl3* mutant allele (Supplementary Fig. 1a), and showed that mutations in *TTL1* and *TTL4*, but not *TTL2*, also caused venation defects that were markedly enhanced in a triple *ttl1 ttl3 tt4* mutant (from now on referred to as *ttl134*) (Supplementary Fig. 1a).

TTL3 has been proposed as an adaptor protein of BRL2 that, through association with other proteins modulate vein formation ^13^ TTL3, as other TTL proteins from other plant species ^11,12^, are characterized by the presence of 6 tetratrico peptide repeats (TPR) and a C-terminal domain with homology to thioredoxins. An *in silico* structural analysis of TTL3 predicts the presence of an intrinsically disordered region (IDR) at the N-terminus (Supplementary Fig. 2) with the rest of the protein forming a horseshoe-shaped structure composed of multiple helix-turn-helix motifs (Fig. 1a). This structure is consistent with TTL3 being involved in protein-protein interactions and the assembly of multi-protein complexes ^20–22^.

**Figure 1.**
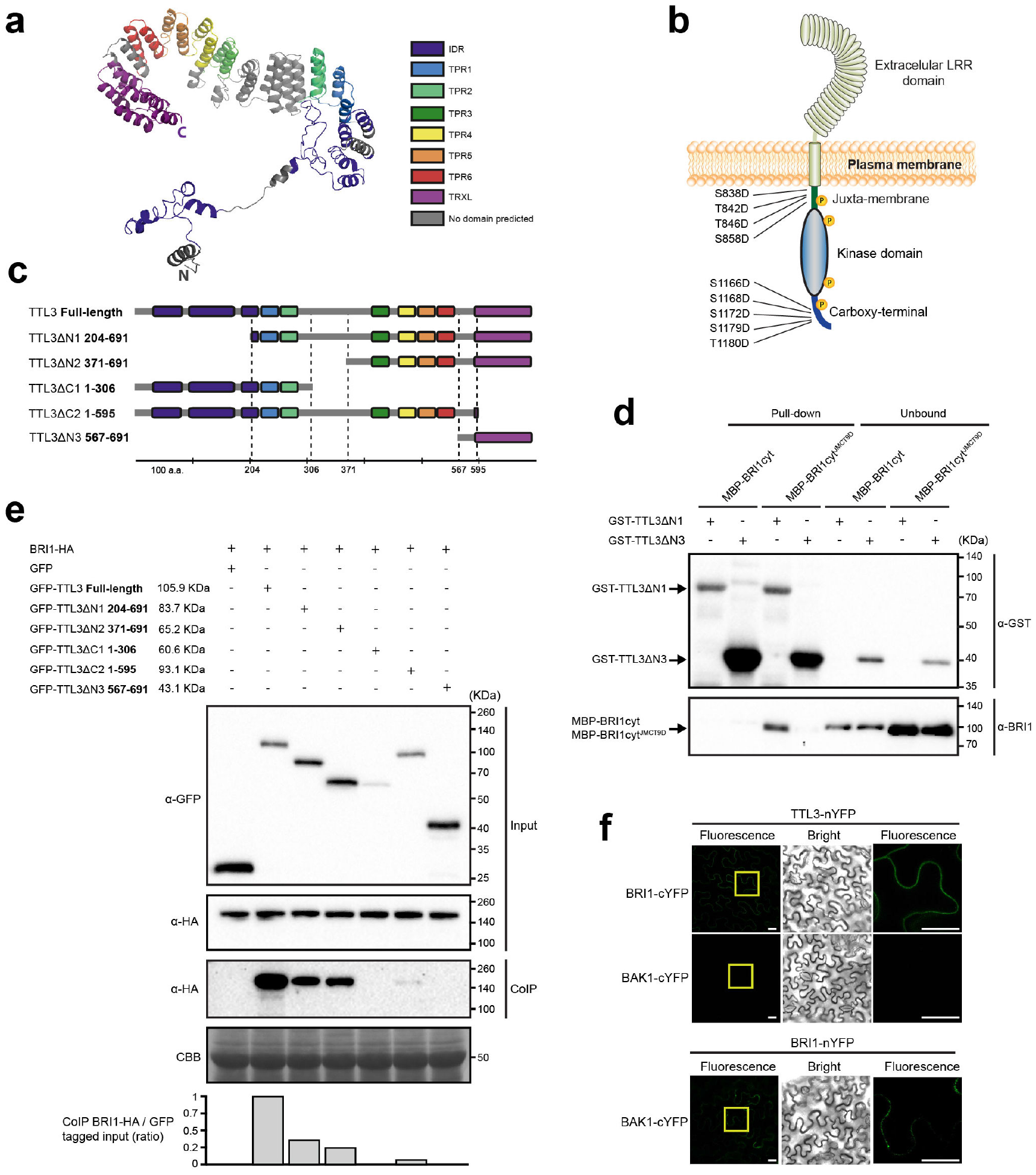
TTL3 interacts with BRI1 *in vivo* and *in vitro.* **a** The structural model of the TTL3 protein predicted *in silico* using I-TASSER server ^69^ and processed by PyMOL (Schrödinger). IDR, interisticaly disorder region; TPR, tetratricopeptide repeats; and TPRX thioredoxin-like domain with homology to thioredoxins; N; N-terminus; C; C-terminus. **b** Schematic representation of BRI1 protein and the nine Serine/Threonine residues of the juxta-membrane and carboxyl-terminal domains that were substituted by Aspartic Acid in the BAK1-independent BRI1-constitutive (phosphomimetic) active form BRI1cyt^JMCT9D 70^. **c** Schematic representations of full-length and different truncated versions of TTL3 protein. Numbers indicates first and last amino acids of TTL3 truncated proteins. IDR, interisticaly disorder region; TPR, tetratricopeptide repeats; TPRX, thioredoxin-like domain with homology to thioredoxins; domains and protein fragments interspacing the conserved domains are represented with the same color code as in **a**. **d** TTL3ΔN1 interacts with BRI1cyt^JMCT9D^ *in vitro*, as shown by GST-pull down assay. GST-TTL3ΔN1 and GST-TTL3ΔN3 were detected with nti-GST antibody. MBP-BRI1cyt and MBP-BRI1cyt^JMCT9D^ were detected using specific anti-BRI1 antibodies ^71^. Pull-down reflects 20% of the total pulled-down proteins. Unbound reflects 1% of the total unbound fraction. **e** BRI1-HA co-immunoprecipitates with GFP-TTL3 full length and GFP-TTL3 truncated versions ΔN1, ΔN2 and ΔC1. Numbers indicate first and last amino acids of TTL3 truncated proteins. BRI1-HA was transiently co-expressed in *N. benthamiana* with GFP-TTL3 full length and truncated versions and GFP tagged protein was immunoprecipitated using anti-GFP Trap beads. Total (input), immunoprecipitated (IP) and Co-Immunoprecipitated (CoIP) proteins were analyzed by western blotting. Equal loading was confirmed by Coomassie blue staining (CBB) of input samples. GFP and HA tagged proteins were detected with anti-GFP and anti-HA antibody, respectively. **f** Bimolecular fluorescent complementation (BiFC) confirms the association of TTL3 with BRI1 but not with BAK1. Leaves of *N. benthamiana* were infiltrated with the *Agrobacterium* strains harboring constructs to express TTL3 and BRI1 proteins fused to the N-terminus of the YFP and, BRI1 and BAK1 proteins fused to the C-terminus of the YFP. Using the same settings in the confocal microscope, YFP fluorescence is observed when TTL3-nYFP is co-expressed with BRI1-cYFP, but no YFP fluorescence is detected when TTL3-nYFP is coexpressed with BAK1-cYFP. A weak YFP fluorescence is observed when BRI1-nYFP is co-expressed with BAK1-cYFP. From left to right columns, images show BiFC YFP fluorescence in green, bright field, and 4× magnification of BiFC YFP fluorescence of the region delimited by the yellow square. Scale bars represent 20 μm. All experiments were repeated at least three times with similar results.

A previous report indicated a role for *TTL3* in BR responses ^13^, and the similarity between BRL2 and BRI1 kinase domains (Supplementary Fig. 3) suggested that TTL3 could also interact with the BRI1 cytoplasmic domain. We therefore tested the *in vitro* direct interaction of TTL3 with the BRI1 cytoplasmic region, which includes the juxta-membrane (JM), the kinase domain and the carboxy-terminal (CT) domain (BRI1cyt) (Fig. 1b). While BRI1cyt was soluble when fused to an MBP tag (Supplementary Fig. 4), we were unable to produce full-length TTL3 protein fused to GST despite many attempts (data not shown) probably due to low stability caused by the IDR ^24^ We could however produce in *E. coli* two different soluble fragments: TTL3 lacking the N-terminus IDR (TTL3ΔN1) and TTL3 containing the TRLX domain (TTL3ΔN3) (Fig. 1c; Supplementary Fig. 4). Using an *in vitro* GST pull-down assay we did not detect interaction of BRI1cyt with either TTL3ΔN1 or TTL3ΔN3 (Fig. 1c, d). Because the activation of BRI1 is dependent on BRI1-ASSOCIATED KINASE 1 (BAK1) transphosphorylation on specific residues at the JM and CT (Wang et, 2008) we used a BAK1-independent BRI1 constitutively-active (phosphomimetic) form BRI1cyt^JMCT9D^ in which nine serines and threonines have been substituted by aspartic acid at the JM and CR domains (Wang et al., 2008) (Fig. 1b). In this case, BRI1cyt^JMCT9D^ was pulled down by TL3ΔN1, but not by TTL3ΔN3 (Fig. 1c, d). This indicates that TTL3 predominantly interacts with active BRI1 form that is independent of BAK1 activation, and that this interaction occurs between the TPR domains, but not the TRLX domain of TTL3.

Next, we investigated this interaction *in vivo* by performing co-immunoprecipitation (Co-IP) assays after transient expression of tagged full-length TTL3 and BRI1 in *Nicotiana benthamiana.* After immunoprecipitation of GFP-TTL3 and free GFP using GFP-Trap beads, we detected a strong specific interaction between GFP-TTL3 and BRI1-HA (Fig. 1e. Lanes 1 and 2). Additional Co-IP experiments using a C-terminally GFP tagged TTL3 protein (TTL3-GFP) co-expressed with BRI1-HA (Supplementary Fig. 5a) and BRI1-GFP co-expressed with TTL3-HA (Supplementary Fig. 5b) further confirmed the specificity of TTL3-BRI1 interaction and indicated that the position and tag used in the Co-IP experiments does not affect their interaction *in planta.*

We further used Co-IP assays to map the interaction domains of TTL3 required for the interaction with BRI1. We performed this analysis *in planta* in order to determine the possible role of the IDR domain in the interaction, which was not possible using *in vitro* assays. We generated a series of truncated TTL3 fragments with deletions at the N-terminus (TTL3ΔN1, TTL3ΔN2, TTL3ΔN3) and at the C-terminus (TTL3ΔC1, TTL3ΔC2), transcriptionally fused to GFP at the N-terminus (Fig. 1c) and co-expressed with BRI1-HA in *N. benthamiana* leaves. Expression analysis of the truncated proteins indicated that all accumulated at the expected molecular size (Fig. 1e, Input). TTL3ΔC1 and TTL3ΔC2 constructs, both lacking the TRLX domain, showed lower accumulation than the other constructs, suggesting that TRLX is important for protein stabilization.

Three of the five truncated TTL3 proteins, *i.e.* GFP-TTL3ΔN1 and GFP-TTL3ΔN2 and GFP-TTL3ΔC2 co-immunoprecipitated BRI1-HA with different efficiency - all having in common TPR3 to TPR6 (Fig. 1c) - indicating that these domains are essential for the interaction, which is consistent with the *in vitro* data (Fig. d). In order to better evaluate the interaction of the different TTL proteins fragments and BRI1, the amount of co-immunoprecipitated BRI1-HA was normalized relative to the amount of protein input (Fig. 1e). The strongest interaction occurs with the full-length TTL3 protein, indicating that all domains contribute to stabilize the interaction with BRI1. A lower but similar interaction was observed with GFP-TTL3ΔN1 and GFP-TTL3ΔN2, both containing the TRLX domain, indicating that this domain is important for a stable interaction although it is not sufficient to interact with BRI1 *in vitro* or *in vivo* (Fig. 1c, e). Consistent with this result, removing the TRLX region in GFP-TTL3ΔC2 greatly reduced the interaction between TTL3 and BRI1 (Fig. 1c, d, e).

Finally, the interaction between BRI1 and TTL3 was also investigated using bimolecular fluorescence complementation (BiFC) assays in *N. benthamiana* leaves, which provide additional information about the subcellular localization of the interaction. As shown in Fig. 1f, co-expression of TTL3-nYFP with BRI1-cYFP or BRI1-nYFP with TTL3-cYFP (Supplementary Fig. 5c) reconstituted functional YFP proteins at the plasma membrane, which confirm the interaction and is consistent with the plasma membrane localization of BRI1.

BAK1, also known as SERK3, and other SERK proteins are transmembrane kinases that function as BR co-receptors ^25^. Similar Co-IP experiments using TTL3-GFP and BAK1 transiently co-expressed in *N. benthamiana* indicated that, contrary to BRI1, TTL3 does not associate *in vivo* with BAK1 (Supplementary Fig. 5d). This result was verified by BiFC assays in *N. benthamiana* leaves. Confocal microscopic analyses revealed that coexpression TTL3-nYFP with BAK1-cYFP (Fig. 1f) and also BAK1-nYFP with TTL3-cYFP (Supplementary Fig. 5c) did not reconstitute functional YFP proteins. To confirm that BiFC BAK1 constructs were functional, we performed BiFC between BRI1 and BAK1 resulting in positive signals (Fig. 1f; Supplementary Fig. 5c)

### *ttl* mutants show defects in BR responses

The interaction of TTL3 with BRI1 supports a role of TTL3 in BR signaling. Quantitative RT-PCR analyses indicate that the expression of *TTL1, TTL3*, and *TTL4* is induced by BR ^14^, which is also supported by available transcriptomic data (Supplementary Fig 6a). This up-regulation of the *TTL* genes in response to BR was confirmed at cellular level by analyzing transgenic plants transformed with the reporter β-glucuronidase gene driven by each of the *TTL* promoters (Supplementary Fig 6b).

Next, we analyzed the sensitivity to exogenous epibrassinolide (eBL) by measuring root growth in the presence or absence of 100 nM eBL in single *ttl* mutants, the triple *ttl134* mutant and *bak1-4*, a well-established mutant affected in BR responses ^26–29^. Single *ttl* mutants, the *ttl134* mutant, and the *bak1-4* mutant showed a similar root growth to Col-0 in control conditions (Fig. 2a; Supplementary Fig. 7a). However, *bak1-4, ttl1, ttl3*, and *ttl4* show increased root length than Col-0 control or *ttl2* in the presence of eBL (Fig. 2a; Supplementary Fig. 7b). This decreased sensitivity to eBL of single *ttl* mutants was strongly enhanced in the *ttl134* (Fig. 2a; Supplementary Fig 7b). Root growth sensitivity to eBL of the *ttl134* mutant was then compared, in addition to *bak1-4*, to well characterized genotypes affected in BR responses such as *serk1-1* and the double *serk1-1 bak1-4* mutant^26^ In control conditions, all genotypes grew similarly, with the exception of *serk1-1 bak1-4*, which showed reduced root growth (Fig. 2b; Supplementary Fig 7c) as previously reported ^30,31^. In the presence of 100 nM eBL, the root growth reduction of the Col-0 control was significantly higher than for the rest of the genotypes, including *ttl134* (Fig. 2b; Supplementary Fig 7d), while *serk1-1 bak1-4* double mutant was almost insensitive to eBL, as it showed a similar root growth in control and eBL-supplemented media (Fig. 2b; Supplementary Fig 7c, d).

**Figure 2.**
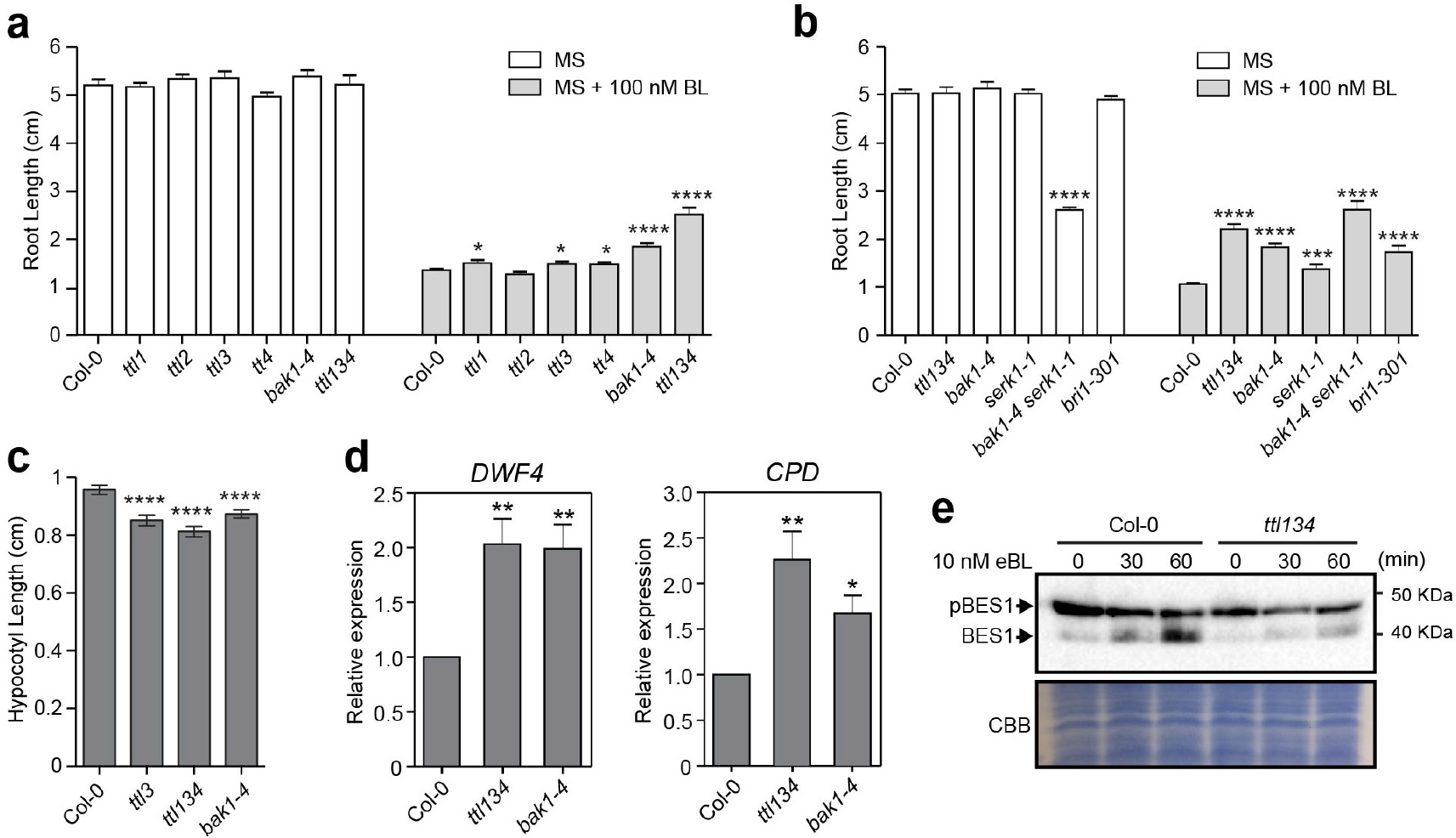
*TTL1, TTL3 and TTL4* genes play a positive role in BR signaling pathway. **a** *ttl1, ttl3, ttl4* and *ttl134* show root growth hyposensitivity to BR. Statistical analysis of root length measurements of Col-0, *ttl*, and *bak1-4* mutants in control conditions (MS) and in response to BL. Seedlings were grown in long days for 4 days in half-strength MS agar solidified medium and then transferred to half-strength MS agar solidified medium (MS) or half-strength MS agar solidified medium supplemented with 100 nM of Brassinolide (MS + 100 nM eBL) and root length was measure 6 days later. Asterisks indicate statistical differences between mutant vs Col-0 determined by the unpaired *t-test* (* P ≤ 0.05, ** P ≤ 0.01, *** P ≤ 0.001 **** P < 0.0001). Data represent mean values, error bars are SEM, n≥35 seedlings per experiment. The experiment was repeated three times with similar results. **b.** Root length responses to eBL of wild-type Col-0, *ttl134* and BR perception mutants. Seedlings were grown and root length was analyzed as described in a. Asterisks indicate statistical differences between mutant vs Col-0 as determined by the unpaired *t-test* (*** P ≤ 0.001 **** P ≤ 0.0001). Data represent mean values, error bars are SEM, n=30 seedlings per experiment. The experiment was repeated three times with similar results. **c** Defective hypocotyl elongation in *ttl* mutants. Col-0, *ttl3, ttl134* and *bak1-4* seedlings were grown for 4 days in long-day photoperiod in half-strength MS agar solidified medium. Seedlings with the same size were then placed in the dark and hypocotyl elongation was measure 3 days later. Asterisks indicate statistically difference significances between Col-0 vs the indicated genotype as determined by the unpaired t-test (**** P ≤ 0.0001), values are mean, error bars are SEM, n = 80 seedlings per experiment. The experiment was repeated twice with similar results. **d** BR-responsive genes *DWF4* and *CPD* show induced expression in *ttl134* and *bak1-4* relative to Col-0 seedlings. Seeds were germinated in half-strength MS agar solidified medium and grown vertically in long-day photoperiod conditions. 5-day-old seedlings were transferred to half-strength MS liquid medium and after 5 days of acclimation, relative expression level of *DWF4* and *CPD* was measured by quantitative reverse transcriptase PCR (qPCR). The expression of *DWF4* and *CPD* was first normalized to the expression of *ACTIN2* gene and represented relative to the expression of Col-0. The data are shown as mean ± SEM from at least three independent biological replicates. Asterisks indicated statistically significant differences between the indicated genotype vs Col-0 as determined by the unpaired t-test (* P ≤ 0.05, ** P ≤ 0.01). The experiment was repeated three times with similar results. **e** Phosphorylation status of BES1 in response to exogenous applied BR in Arabidopsis Col-0 and *ttl134.* Ten-day-old seedlings pre-treated for 3 days with the BR biosynthetic inhibitor brassinazole (BRZ) to deplete the endogenous pool of BRs were submitted to 10 nM eBL treatment for 0, 30 and 60 minutes. Total proteins were analyzed by an immuoblotting assay with a specific anti-BES1 antibody ^72^. The upper band corresponds to phosphorylated BES1 (pBES1) and the lower one to dephosphorylated BES1 (BES1). The experiment was repeated two times with similar results.

Hypocotyl elongation in the dark is dependent on active BR signaling ^32^ We analyzed hypocotyl elongation in the dark of *ttl3, ttl134* and *bak1-4* as a read-out of defective BR signaling ^33^. As previously reported, *bak1-4* showed a reduction in hypocotyl elongation relative to Col-0 ^34,35^(Fig. 2c; Supplementary Fig 8). Similar to *bak1-4, ttl3* and *ttl134* mutants presented shorter hypocotyls than Col-0 (Fig. 2c; Supplementary Fig 8).

To investigate the contribution of *TTL* genes to BR responses at the molecular level, we first studied the expression of the BR-regulated genes *CPD1* and *DWF4* in Col-0, *bak1-4*, and the triple *ttl134* mutants. As shown in Fig. 2d, *DWF4* and *CDP1* expression was around two-fold higher in *ttl134* and *bak1-4* compared to the Col-0 control. This increased *CPD1* and *DWF4* expression has been reported for BR signaling mutants such as *bri1-5* ^36^, *bri1-301* ^26^ and *bik1* ^37^, and is caused by a lack of feedback regulation in the expression of these biosynthetic genes ^38–40^. Second, we investigated the phosphorylation status of BES1 in Col-0 and the *ttl134* mutant in response to eBL. Because the BR biosynthetic genes *DWF4* and *CDP1* are induced in *ttl134*, and to fully capture the BR signaling capacity of *ttl134*, we first pretreated the seedlings with BR biosynthesis inhibitor brassinazole (BRZ). Without BR treatment, a strong phosphorylated BES1 (pBES1) band and a weak unphosphorylated (BES1) band are present in Col-0 and *ttl134* (Fig. 2e). As expected, BR treatment caused an increase of dephosphorylated BES1 in Col-0 due to activation of the pathway. However, eBL caused little dephosphorylation of pBES1 in *ttl134* seedlings (Fig. 2e), confirming a defective BR signaling in *ttl134*.

### BRs regulate the cytoplasmic/plasma membrane localization of TTL3

To further explore how TTL3 functions in BR signaling we analyzed its subcellular localization. Although the BiFC interaction of TTL3 with BRI1 suggests a plasma membrane localization of TTL3, expression of a C-terminal GFP-tagged TTL3 in *N. benthamiana* indicated a predominant cytoplasmic localization in basal conditions (Supplementary Fig. 9a). However, plasmolyzed cells show the presence of GFP-TTL3 in Hechtian strands, indicative that TTL3 also associated with the plasma membrane (Supplementary Fig. 9b). In order to gain further insight into TTL3 localization, a genomic fragment including a 1.7 kb *TTL3* promoter region upstream of the start codon was transcriptionally fused to GFP to generate the *TTL3p::TTL3g-GFP* construct and transformed into *ttl3* and *ttl134* mutants using *A. tumefaciens.* After confocal analysis of a large number of independent stable transgenic lines, we selected two homozygous lines, one in *ttl3* background (hereafter referred to as *TTL3-GFP 1.2*) and another in *ttl134* background (*TTL3-GFP 2.4)*, which presented noticeable fluorescence signals. Venation defects of *ttl3* and *ttl134* were restored to levels similar to Col-0 in *TTL3-GFP 1.2* and *TTL3-GFP 2.4*, (Supplementary Fig.1b). Furthermore, root growth of *TTL3-GFP 1.2* (Supplementary Fig.10a, b) and *TTL3-GFP 2.4* (Fig. 3a, b, c) were restored to wild type levels in the presence of eBL, indicative of a functional TTL3-GFP protein.

**Figure 3.**
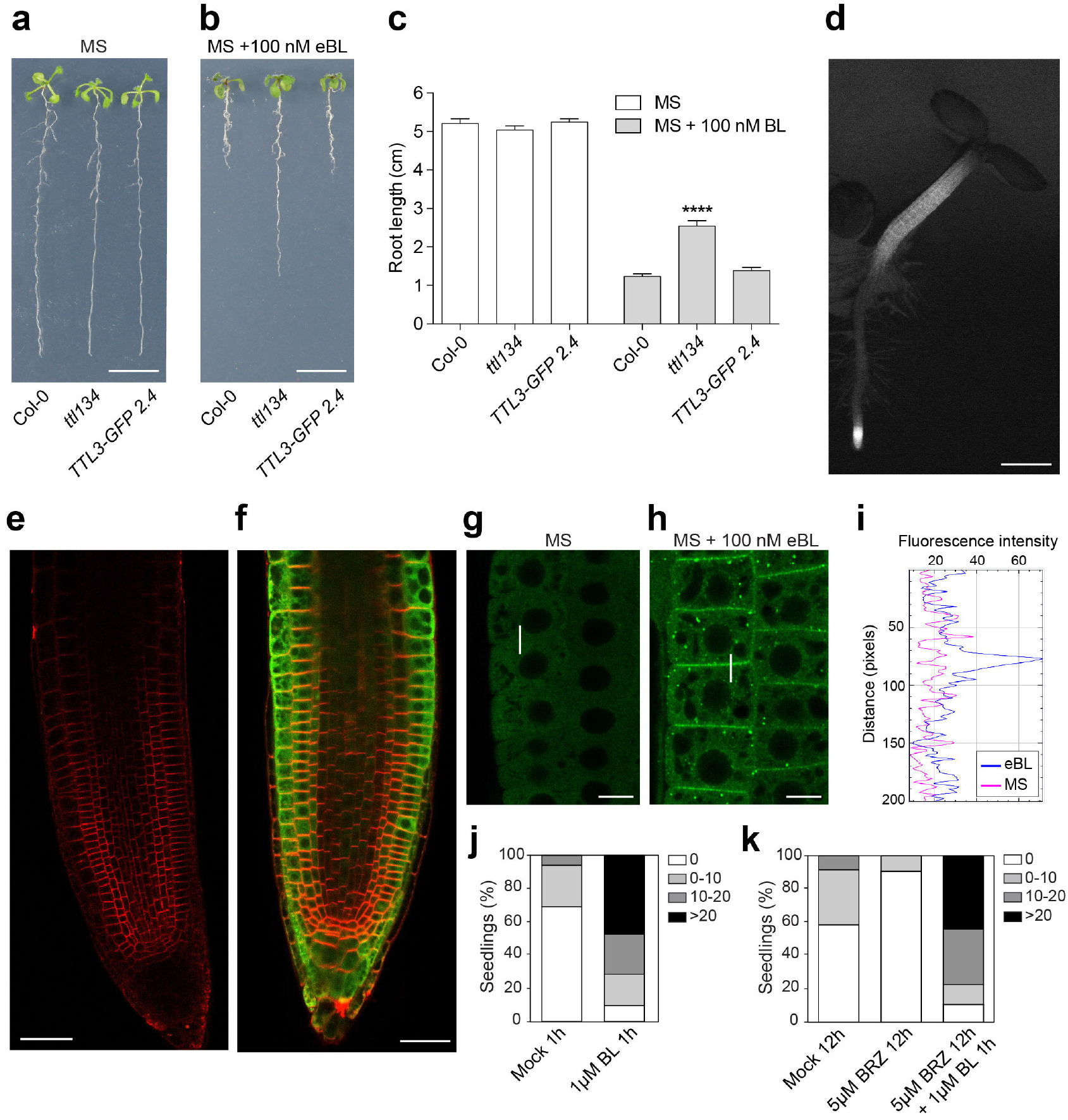
BRs regulate the cytoplasmic/plasma membrane localization of TTL3. **a-c** The root growth responses to eBL of the *ttl134* triple mutant are complemented in the *TTL3-GFP 2.4.* Seedlings were grown for 4 days in halfstrength MS agar solidified medium and then transferred to half-strength MS agar solidified medium (**a**) or half-strength MS agar solidified medium supplemented with 100 nM of Brassinolide (**b**). **a** Representative picture of seedlings 6 days after treatment. Scale bar represents 1 cm. c Statistical analysis of root length of Col-0, *ttl134* and the complementation line *TTL3-GFP* 2.4. Asterisks indicate statistically significant differences between the indicated genotype vs Col-0 as determined by the unpaired *t-test* (**** P ≤ 0.0001). Data represent mean values, error bars are SEM, n=30 seedlings per experiment. The experiment was repeated three times with similar results. **d** Expression pattern of *TTL3-GFP* in 3-day-old *TTL3-GFP 2.4* Arabidopsis seedlings. Image was captured using conventional wide field fluorescence microscopy with a GFP filter. Scale bar represents 500 μm. **e-f** Longitudinal median section of root tips of a 3-day-old Col-0 (**e**) and *TTL3-GFP 2.4* as observed by laser scanning confocal microscopy (**f**). Images show a merge of green channel showing TTL3-GFP expression and red channel showing plasma membrane stained with FM4-64. Scale bar represents 20 μm. **g-i** Confocal images showing localization of TTL3-GFP in epidermal cells from root meristematic zone in 4-day-old Arabidopsis *TTL3-GFP 2.4* in half-strength MS agar solidified medium, in control conditions (1 hour treatment with eBL solvent) (**g**) or after 1 hour of 1 μM eBL treatment (**h**) in half-strength MS agar liquid medium. Scale bar represents 10 μm (horizontal bar). **i** Quantification of fluorescent protein signal in plasma membrane vs cytoplasm. Line scan measurements spanning membrane and cytoplasm were carried out (represented in **g** and **h** as a vertical white line), and representative plot profiles of sample measurements are presented. **j-k** Quantification of the cytoplasmic and PM localization of TTL3-GFP in 4-day-old Arabidopsis *TTL3-GFP 2.4* seedlings treated for 1 hour with 1 μM eBL (**j**), or pre-treated for 12 hour with 5 μM BRZ prior to 1 μM eBL application for 1 hour (**k**). Analyses were carried out counting the number of cells with dual cytoplasmic/plasma membrane localization in meristematic and transition zone for each analyzed root using confocal microscopy. Seedlings were grouped in categories according to the number of cells that presented this dual localization, and the percentage of seedlings displaying each category depicted at right side panel was calculated. Represented categories (right side panel) indicate the number of cells per seedling with dual cytoplasmic/plasma membrane localization. At least 16 seedlings per treatment, and approximately 200 cells from epidermis, cortex and endodermis per seedling of the meristematic region of the root tip were analyzed.

We then used *TTL3-GFP 2.4* (which showed a stronger fluorescence signal than *1.2*) to analyze the cellular and subcellular localization of TTL3. Examination under a stereomicroscope indicated that TTL3-GFP accumulated mainly at the root tip and the hypocotyl of Arabidopsis seedlings (Fig. 3d). This accumulation coincides with cells that undergo strong BR signaling leading to active growth, and highly resembles the accumulation pattern of BRI1-GFP ^41–43^. Cellular analysis using confocal microscopy was performed in 3-day-old roots, simultaneously localizing TTL3-GFP with the FM4-64, a lipophilic red dye that labels the plasma membrane and tracks plasma membrane-derived endosomes ^44^ In Col-0 control roots, no GFP signal was detected (Fig. 3e), while analysis of *TTL3-GFP 2.4* revealed the presence of TTL3-GFP in all cell files of the root apical meristem (Fig. 3f). Further up, in the meristematic region, TTL3-GFP showed a predominant localization in the outer cell layers (epidermis and cortex) (Fig. 3f).

At the subcellular level, TTL3-GFP mostly showed a cytoplasmic localization in the root meristematic cells (Fig. 3g). However, we sometimes observed seedlings that, in addition to the cytoplasmic GFP localization, showed GFP signal at the plasma membrane. Therefore, we quantified the plasma membrane localization of TTL3-GFP (see Figure legend and Methods section for details) in control conditions and found that in ~30% of the seedlings some cells showed plasma membrane localization of TTL3-GFP (Fig. 3j).

Interestingly, treatment with 1 μM eBL, a concentration previously used to analyze short-term BKI1 dynamics ^45^, increased the amount of TTL3-GFP protein (Supplementary Fig. 11) and caused a relocalization of TTL3-GFP from the cytoplasm to the plasma membrane (Fig. 3h, j; Supplementary Fig. 12a, b). A detailed quantification indicated that eBL treatment cause a drastic increase in the amount of seedlings and the number of cells per seedling with plasma membrane-localized TTL3-GFP (Fig. 3j). eBL treatment also caused the appearance of GFP-labeled intracellular structures (Fig. 3h), although these intracellular TTL3-GFP structures do not colocalize with FM4-64 (Supplementary Fig. 12), discarding the possibility that they may correspond to plasma membrane-derived endosomes, and thus their identity remains elusive.

Consistent with the possibility that the plasma membrane localization of TTL3-GFP in seedlings grown in control medium was caused by endogenous BRs, the percentage of seedlings with plasma membrane signal decreased from ~30% to ~5% after treatment with BRZ (Fig. 3k). Further treatment of these seedlings with eBL reverted this effect and increased the plasma membrane localization of TTL3-GFP (Fig. 3k).

### TTL3 associates with the BR signaling components BSK1, BSU1 and BIN2 and directly interacts with BSU1

Our previous analyses indicate that TTL3 is involved in BR signaling probably through the scaffolding of BR signaling components. Using Co-IP and BiFC in *N. benthamiana* we investigated the possible association of TTL3 with other core components of BR signaling. TTL3 strongly associates with BSK1 in both Co-IP and BiFC assays (Fig. 4a, b). BiFC between TTL3 and BSK1 was also obtained when we exchanged nYFP and cYFP tags (Supplementary Fig. 13) and consistent with the plasma membrane localization of BSK1, the BiFC signal for BSK1-TTL3 was observed at the plasma membrane. TTL3 also associates with BSU1 and BIN2 in both Co-IP and BiFC assays (Fig. 4b, c, d). Although BSU1 and BIN2 present a dual nuclear and cytoplasmic localization ^46,47^, BiFC signals were only observed in the cytoplasm for both TTL3-BSU1 and TTL3-BIN2, which is consistent with the lack of TTL3 protein in the nucleus (Fig. 4b). A cytoplasmic BiFC signal was also obtained when YFP halves were interchanged among TTL3-BSU1 and TTL3-BIN2 (Supplementary Fig. 13). Two BSU1 bands with different mobility in SDS-polyacrylamide gel electrophoresis were obtained after expression in *N. benthamiana.* This apparent difference in size is likely caused by a different phosphorylation status (Fig. 4c), and interestingly, TTL3 mainly associated with the faster mobility BSU1 band (Fig. 4c). Reducing endogenous BRs by BRZ treatment decreased the relative amount of the lower band (Supplementary Fig. 14), suggesting that this band corresponds to the active (dephosphorylated) BSU1 form.

**Figure 4.**
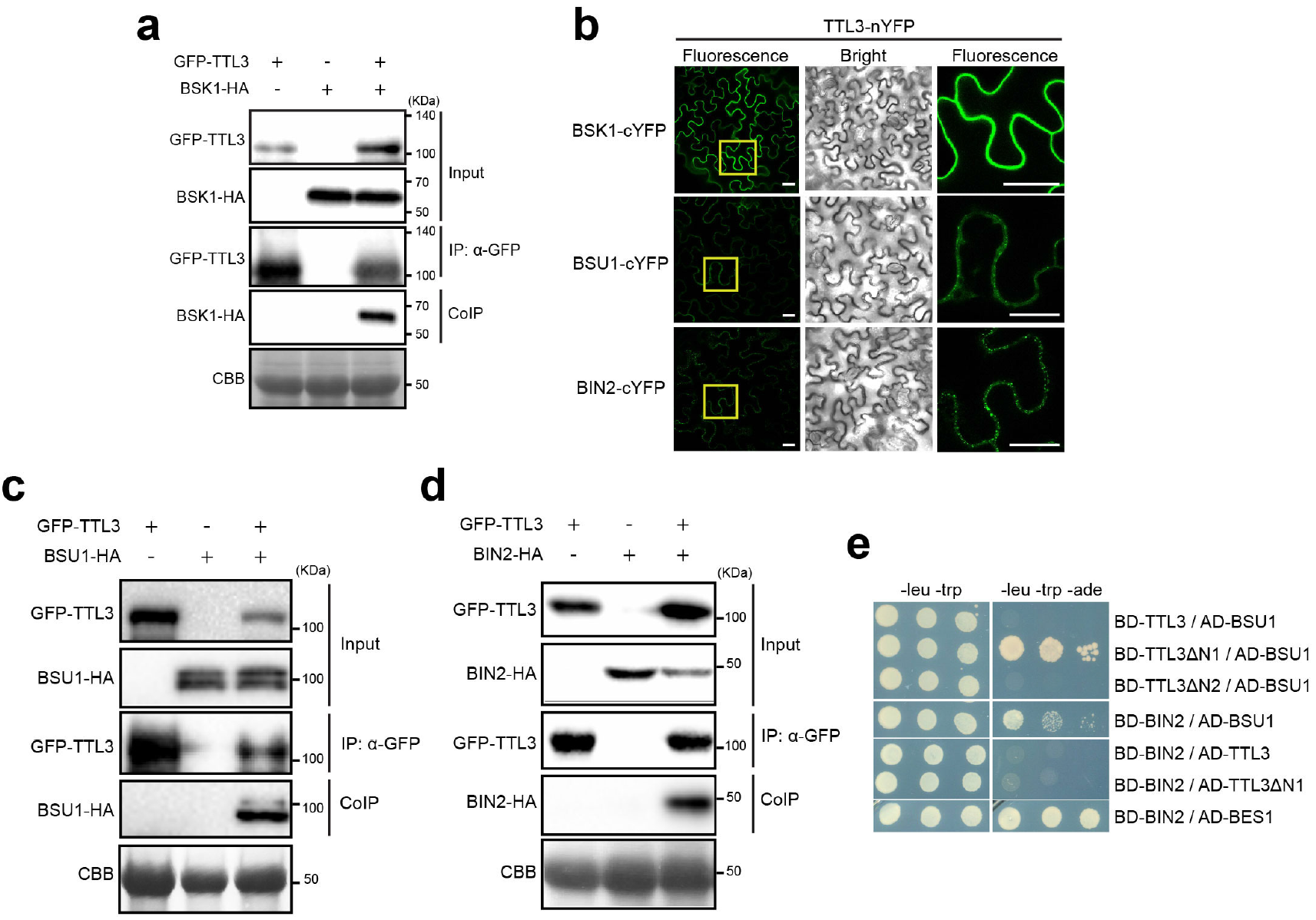
TTL3 associates with BSK1 and BIN2 and directly interacts with BSU1. **a** BSK1 co-immunoprecipitates with TTL3. BSK1-HA and GFP-TTL3 were transiently expressed in *N. benthamiana.* GFP-TTL3 was immunoprecipitated with anti-GFP Trap beads. Total (input), immunoprecipitated (IP) and Co-Immunoprecipitated (CoIP) proteins were analyzed by western blotting. Equal loading was confirmed by Coomassie blue staining (CBB) of input samples. GFP-TTL3 and BSK1-HA were detected with anti-GFP and anti-HA antibody, respectively. **b** BiFC assays confirm the association of TTL3 with BSK1, BSU1 and BIN2. Leaves of *N. benthamiana* were agroinfiltrated with the *Agrobacterium* strains harboring a construct to express TTL3 protein fused to the N-terminus half of the YFP and BSK1, BSU1 and BIN2 protein fused to the C-terminus half of the YFP. Using the same settings in the confocal microscope, YFP fluorescence is observed when TTL3-nYFP is co-expressed with BSK1-cYFP, BSU1-cYFP or BIN2-cYFP. From left to right columns, images show BiFC YFP fluorescence in green, bright field, and 4× magnification of BiFC YFP fluorescence of the region delimited by the yellow square. Scale bars represent 20 μm. **c** BSU1 co-immunoprecipitates with TTL3. GFP-TTL3 and BSU-HA proteins were transiently expressed in *N. benthamiana*, immunoprecipitated and analyzed as described in **a**. GFP-TTL3 and BSU1-HA were detected with anti-GFP and anti-HA antibodies, respectively. **d** Yeast-two-hybrid assays to determine the interaction of full-length TTL3, the TTL3 fragment TTL3ΔN1 (amino acid 204-691) and the TTL3 fragment TTL3ΔN2 (amino acid 371-691) with BIN2 and BSU1. Growth on plasmid-selective media (left column) and interaction-selective media (lacking adenine, right column) are shown. **e** BIN2 co-immunoprecipitates with TTL3. BIN2-HA and GFP-TTL3 proteins were expressed in *N. benthamiana*, immunoprecipitated and analyzed as described in **a**. GFP-TTL3 and BSU1-HA were detected with anti-GFP and antiHA, respectively.

Next, we investigated possible direct interactions between TTL3 and the cytoplasmic BR signaling components BSU1 and BIN2, using yeast two-hybrid assays. Using a full-length TTL3 protein, we did not find interactions with any of the investigated BR components, despite obtaining previously positive reported interactions such as BIN2 with BSU1 and with BES1 (Fig. 4e). Western blot analysis indicated that BD-TTL3 fusion protein was not detected (Supplementary Fig. 15), similar to what previously occurred in *E. coli.* Therefore, we generated additional yeast two-hybrid constructs using the TTL3ΔN1 and TTL3ΔN2 fragments (Fig. 1c). As shown in Fig. 4e, TTL3ΔN1 but not TTL3ΔN2 interacted with BSU1, indicating that the six TPR domains are required for the interaction. In contrast to BSU1, BIN2 did not interact with TTL3ΔN1 (Fig. 4e), despite the positive interactions of BIN2 with BSU1 or BES1 were detected (Fig. 4e). These data indicate that the six TPR of TTL3 are required for the *in vitro* interaction with BSU1 while *in vivo* data suggests that TTL3 preferentially associates with the active (phosphorylated) BSU1.

### TTL3 interacts with the transcription factors BZR1 and BES1 and affects BZR1 cytoplasmic/nuclear localization

In the absence of BRs, BIN2 phosphorylates and inactivates BZR1 and BES1, which are the two major transcription factors mediating BR-induced transcriptional changes ^5^. TTL3 associates with BZR1 in Co-IP experiments in *N. benthamiana* (Fig. 5a) and in Arabidopsis mesophyll protoplasts (Fig. 5b). Phosphorylated and dephosphorylated BZR1 and BES1 proteins show a marked difference in mobility in SDS-PAGE upon expression in *N. benthamiana*, (Fig. 5a, b, Supplementary Fig. 16a, b) or in Arabidopsis protoplasts (Fig. 5b) ^48,49^. Interestingly, only the phosphorylated BZR1 (pBZR1) was co-immunoprecipitated with TTL3 (Fig. 5b, Supplementary Fig. 16a) indicating a preferential association of TTL3 with pBZR1. Similarly, TTL3 only co-inmunoprecipitated the phosphorylated BES1 (pBES1) (Supplementary Fig. 16b). BiFC assays further confirmed the *in vivo* association of BZR1 and BES1 with TTL3 (Fig. 4c, Supplementary Fig. 13). While the BiFC signal of TTL3 with plasma membrane BR components results in a smooth YFP fluorescence signal at the plasma membrane (Fig. 1f, Supplementary Fig. 5c, Fig. 4b, Supplementary Fig. 13), the BiFC signal of TTL3 with the cytoplasmic components appear punctated (Fig. 4b, Supplementary Fig. 13, Fig. 5c). A similar punctate BiFC signal has been previously reported for BZR1 with BRZ-SENSITIVE-SHORT HYPOCOTYL1 (BSS1) ^50^ or BES1 with DOMINANT SUPPRESSOR OF KAR 2 (DSK2) ^7^, although its significance remains unknown.

**Figure 5.**
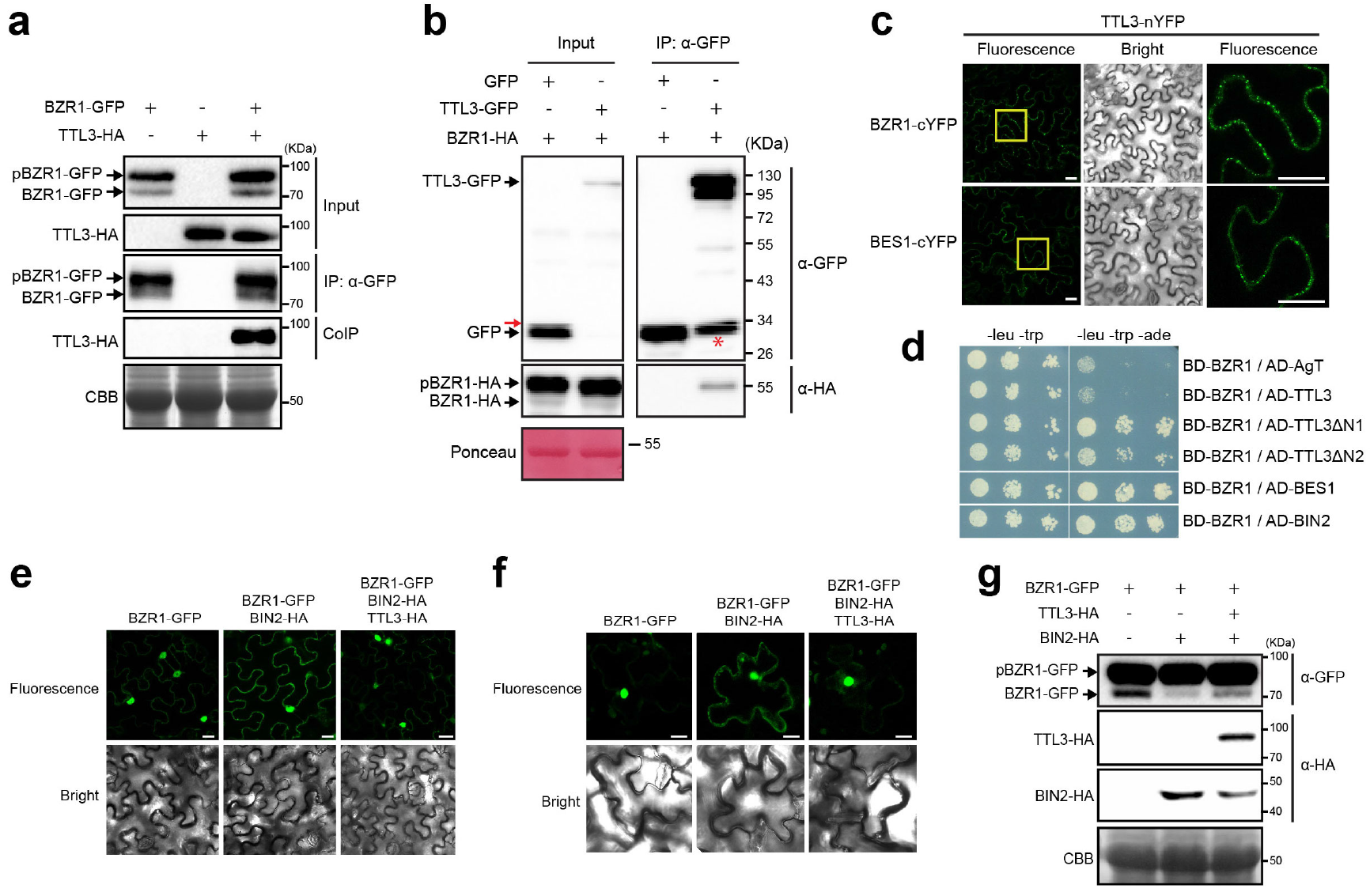
TTL3 interacts with BZR1 and regulates its cytoplasmic/nuclear localization. **a** TTL3 co-immunoprecipitates with BZR1. TTL3-HA and BZR1-GFP were transiently expressed in *N. benthamiana.* BZR1-GFP was immunoprecipitated with anti-GFP Trap beads. Total (input), immunoprecipitated (IP) and Co-Immunoprecipitated (CoIP) proteins were analyzed by western blotting. Equal loading was confirmed by Coomassie blue staining (CBB) of input samples. BZR1-GFP and TTL3-HA were detected with anti-GFP and anti-HA, respectively. The upper band corresponds to phosphorylated BZR1 (pBZR1-GFP) and the lower one to dephosphorylated BZR1 (BZR1-GFP). **b** Co-immunoprecipitation of BZR1-HA with TTL3-GFP expressed in transfected *Arabidopsis* Col-0 protoplasts. Samples were analyzed as in a. Protoplasts cotransfected with free GFP and BRI1-HA, were used as a negative control for Co-IP. Equal loading was confirmed by Ponceau staining of input samples. TTL3-GFP and free GFP were detected with anti-GFP antibody and BRI1-HA was detected with anti-HA antibody. Asterisk indicates GFP that results from proteolytic cleavage of TTL3-GFP. Red arrow indicates an artefact from imaging blot with high sensitivity using Azure c300 Chemiluminescent Western Blot Imaging System **c** BiFC confirms the association between TTL3 and BZR1. Leaves of *N. benthamiana* were agroinfiltrated with the *Agrobacterium* strain harboring a construct to express the TTL3 protein fused to the N-terminus half of the YFP and the BZR1 or BES1 protein fused to the C-terminus half of the YFP. YFP fluorescence is observed when TTL3-nYFP is co-expressed with BZR1-cYFP or BES1-cYFP using confocal microscopy. From left to right columns, images show BiFC YFP fluorescence in green, bright field, and 4× magnification of BiFC YFP fluorescence of region delimited by the yellow square. Scale bars represent 20 μm. **d** Yeast-two-hybrid assays to determine the interaction of BZR1 with TTL3, the TTL3 fragment TTL3ΔN1 (amino acid 204-691), the TTL3 fragment TTL3ΔN2 (amino acid 371-691), BES1 and BIN2. Interaction of BZR1 with a fragment of SV40 large T-antigen (AD-AgT) was also included to show BD-BZR1 selfactivation capacity. Growth on plasmid-selective media (left column) and interaction-selective media (lacking adenine, right column) are shown. **e-f** TTL3 abolishes the cytoplasmic retention of BZR1 by BIN2. Subcellular localization of BZR1-GFP alone, co-expressed with BIN2-HA, and with BIN2-HA and TTL3-HA in *N. benthamiana* leaves (**e**) and in NahG-Arabidopsis leaves (**f**). Images of the GFP signal were obtained by using laser scanning confocal microscopy. Images show a single equatorial plane in *N. benthamiana* leaves (**e**), and a maximum Z-projection of seven 1 μm spaced focal planes from the cell equatorial plane to the cell surface in NahG-Arabidopsis leaves (**f**). Scale bars represent 20 μm. **g** Western blot analysis of the BZR1-GFP proteins transiently expressed alone, co-expressed with BIN2-HA, and co-expressed with BIN2-HA and TTL3-HA in *N. benthamiana* leaves observed by confocal microscopy in e. Proteins were analyzed by western blotting. Equal loading was confirmed by Coomassie blue staining (CBB) of input samples. BZR1-GFP was detected with anti-GFP antibody, while TTL3-HA and BIN2-HA were detected with anti-HA antibody. In the anti-GFP blot, the upper band corresponds to phosphorylated BZR1 (pBZR1-GFP) and the lower one to dephosphorylated BZR1 (BZR1-GFP).

Next we performed a yeast two-hybrid assay between TTL3 and the transcription factor BZR1. As expected (see Fig. 4e), a full-length TTL3 protein did not interact with BZR1, despite detecting the previously described positive interaction between BZR1 and BIN2 ^51^ (Fig. 5d). However we could detect the direct interaction between TTL3ΔN1 (Fig. 1c) and BZR1 (Fig. 5d) and contrary to BSU1, TTL3ΔN2 (Fig. 1c) also interacted with BZR1 (Fig. 5d) indicating that the TPR3 to TPR6 region is sufficient for the TTL3-BZR1 interaction (Fig. 5d).

We next analyzed the effect of TTL3 on the nuclear and cytoplasmic localization of BZR1-GFP. As previously reported, BZR1-GFP in *N. benthamiana* is mainly localized in the nucleus (Fig. 5e), while co-expression of BIN2 together with BZR1-GFP promotes its phosphorylation and its cytoplasmic retention (Fig. 5e) ^52^. Co-expressing TTL3-HA with BZR1-GFP and BIN2-HA suppressed the cytoplasmic retention of BZR1-GFP promoted by BIN2 (Fig. 5e). We also used Arabidopsis plants expressing the salicylate hydroxylase (*NahG*) gene, as these plants are efficiently transiently transformed using *A. tumefaciens* ^53^. Similar to *N. benthamiana*, coexpressing BIN2-HA together with BZR1-GFP increased its cytoplasmic accumulation, which was further abolished by TTL3-HA (Fig. 5f). This BZR1 nuclear/cytoplasmic localization correlates with the dephosphorylation status of BZR1 (Fig. 5g), indicating that TTL3 negatively regulates BIN2 phosphorylation of BZR1 and regulates its activity.

### TTL3 acts as a scaffold by enhancing BZR1-BSK1 interaction

Next, we investigated a possible scaffold function of TTL3 in BR signaling and investigated whether TTL3 affects the association of the plasma membrane-localized BSK1 with cytoplasmic components of BR signaling using BiFC. As shown in Fig. 6a, strong BiFC signal was obtained for BSK1 with BRI1, BSU1 and BIN2 while a weak signal was obtained for BSK1 with BZR1. The strong BiFC signal detected for BSK1 with BRI1 and with BSU1 is expected since this BR signaling components direct interact with BSK1 ^52^. BIN2, although mainly localizes at the nucleus and cytosol, also localizes at the plasma membrane ^47^ and direct interaction with several plasma membrane-localized BSKs in yeast two-hybrid assays was previously reported ^54^ The weak BSK1-BZR1 association is consistent with a previous proteomic study that identified BSK1 as an interactor of BZR1 ^55^. Importantly, when we co-expressed TTL3-HA together with BSK1-nYFP and BZR1-cYFP the BiFC signal was strongly enhanced (Fig. 6b) indicating that TTL3 increases the association between BSK1 and BZR1 at the plasma membrane. Further Co-IP experiments also showed that the amount of BSK1-HA that was immunoprecipitated with BZR1 - GFP was also enhanced upon co-expression of TTL3-mCherry (Fig. 6c). This result strongly supports a scaffolding role of TTL3 that would help bringing a cytoplasmic component such BZR1 with BR signaling components at the plasma membrane such BSK1.

**Figure 6.**
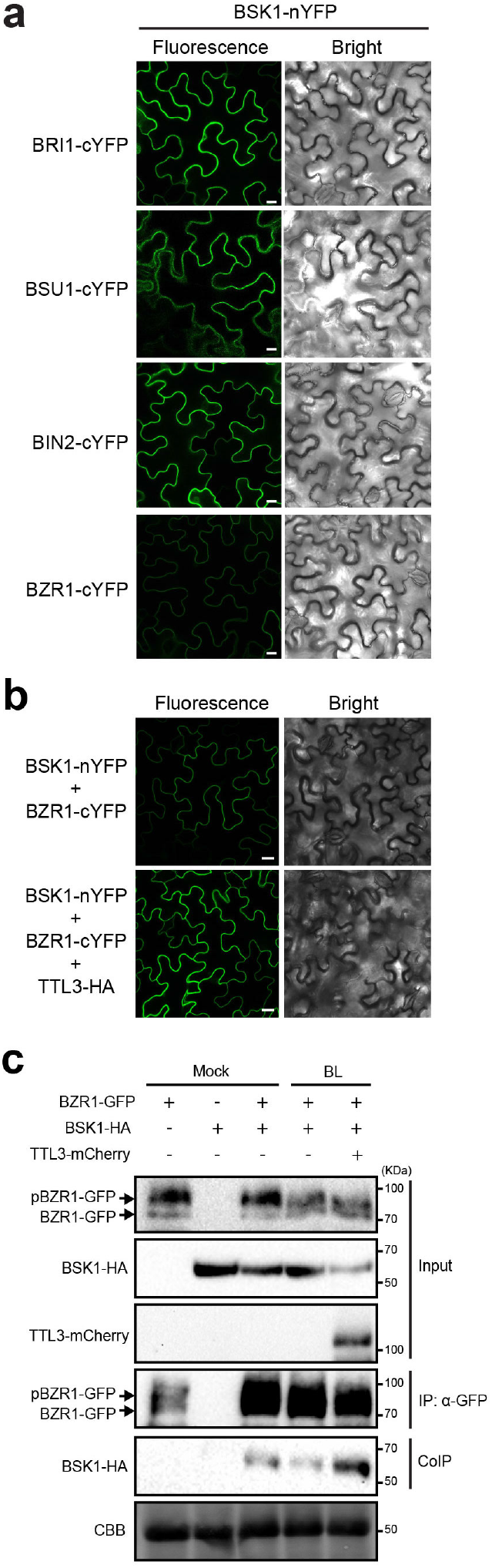
TTL3 acts as a scaffold by enhancing pBZR1-BSK1 interaction. **a** BiFC shows strong association of BSK1 with BRI1, BSU1, BIN2 and weak association with BZR1. *N. benthamiana* leaves were co-agroinfiltrated with the Agrobacterium strains harboring a construct to express the BSK1 protein fused to the N-terminus half of the YFP and the BRI1, BSU1, BIN2 or BZR1 proteins fused to the C-terminus half of the YFP and observed under the laser scanning confocal microscope. Strong fluorescence signals are observed when BSK1-nYFP is co-expressed with BRI1-cYFP, BSU1-cYFP or BIN2-cYFP. Faint YFP signal is observed when BSK1-nYFP is co-expressed with BZR1-cYFP. From left to right columns, images show BiFC YFP fluorescence in green and bright field. Scale bars represent 20 μm. **b** Expression of TTL3 increases the weak BiFC association of BSK1 and BZR1. *N. benthamiana* leaves were co-agroinfiltrated with the *Agrobacterium* strains harboring the corresponding constructs to express the BSK1 protein fused to the N-terminus half of the YFP and the BZR1 protein fused to the C-terminus half of the YFP. *N. benthamiana* leaves were pre-treated with 5 BL for 3 hours before confocal imaging analysis. Co-expression of TTL3-HA together with BSK1-nYFP and BZR1-cYFP highly enhances GFP signal. From left to right columns, images show BiFC YFP fluorescence in green and bright field. Scale bars represent 20 μm. **c** TTL3 increases the amount of BSK1 immunoprecipitated by BZR1. Tagged BSK1-HA and BZR1-GFP proteins were transiently expressed in *N. benthamiana.* BZR1-GFP and BSK1-HA were co-expressed with or without TTL3-mCherry in *N. benthamiana* leaves and were pre-treated with mock or 5 BL for 3 hours as indicated in the figure. BZR1-GFP was immunoprecipitated with anti-GFP Trap beads. Total (input), immunoprecipitated (IP) and Co-Immunoprecipitated (CoIP) proteins were analyzed by western blotting. Equal loading was confirmed by Coomassie blue staining (CBB) of input samples. Co-expression of TTL3-mCherry enhanced the amount of BSK1-HA that CoIP with BZR1-GFP. BZR1-GFP and BSK1-HA were detected with anti-GFP and anti-HA antibody, respectively.

## DISCUSSION

Our study reveals that plant-specific TTL proteins function as positive regulators of BR signaling. The expression of *TTL* genes is induced by BRs and TTL3 shows its highest expression at the root elongation zone and at the hypocotyl, which are areas of high BR activity ^32,56^ A functional TTL3-GFP is mainly localized in the cytoplasm but also shows plasma membrane localization dependent on BR concentration. The *ttl134* mutant is hyposensitive to BR in root growth assays, shows reduced hypocotyl elongation under darkness, has increased expression of BR marker genes *CPD1* and *DWF4* in normal growth conditions, and exhibits reduced BES1 dephosphorylation levels after BR treatment. Furthermore, co-expression of TTL3 together with BZR1 and BIN2 abolishes the BIN2-directed BZR1 cytoplasmic retention in Arabidopsis and *N. benthamiana.* Thus, TTL3 negatively regulates BIN2-phosphorylation and subcellular localization of BZR1 ^48,49,52,57^.

TTL3 protein associates *in vivo* with all core BR signaling components, with the exception of BAK1, and shows direct interaction with BRI1, BSU1 and BZR1. TTL3 contains several defined domains: an IDR at the N-terminus followed by 6 TPR domains involved in protein-protein interactions and assembly of multiprotein complexes, and a region with homology to thioredoxins at the C-terminus. With the exception of the IDR, most of the protein is predicted to form helix-turn-helix. Mapping the interaction domains of TTL3 with BRI1 indicates that the last four TPRs are essential for this interaction, the TRLX domain is important for protein stabilization, and that both TRLX and the IDR contribute to strengthen the interaction. The presence of an IDR in TTL proteins can provide additional advantages in their scaffolding and regulatory function. It was previously reported that IDRs allow their interaction with a large number of interaction partners due to their ability to adopt different conformations thus allowing the assembly of multiple proteins ^58^. We also found that interaction of TTL3 with BSU1 requires all 6 TPRs while only the last four TPR domains are required for the interaction with BZR1.

The BR-related phenotypes, together with the structure of TTL3 and the interactions here described, led us to propose a model in which TTL3 (and probably other TTLs) functions as a scaffold for BR signaling components (Fig. 7). In the absence of BR, TTL3 is localized in the cytoplasm where it forms a complex with phosphorylated BZR1 and BIN2. In these conditions BZR1 is continuously phosphorylated by BIN2, keeping it inactive. Upon BR perception, the activation of BRI1 by BAK1 causes the re-localization of TTL3-GFP to the plasma membrane, which in turn, brings the TTL3-associated BR cytoplasmic components to the plasma membrane causing the assembly of the pathway components (Fig. 7). The small amount of plasma membrane-localized TTL3 in control conditions probably reflects basal BRI1 signaling induced by endogenous BR, as demonstrated by the reduced plasma membrane localization of TTL3 after BRZ treatment (Fig. 3k). The TTL3-dependent assembly of cytoplasmic BR components at the plasma membrane would then promote the inactivation of BIN2 by the BSU1 phosphatase ^52^ Inactivation of BIN2 by active BSU1 (which is preferentially bound by TTL3) will, in turn, cause the dephosphorylation of BZR1 by PP2A. Because dephosphorylated BZR1 do not interact with TTL3, it will be released from the complex and subsequent activation of BR dependent genes in the nucleus will take place (Fig. 7).

**Figure 7.**
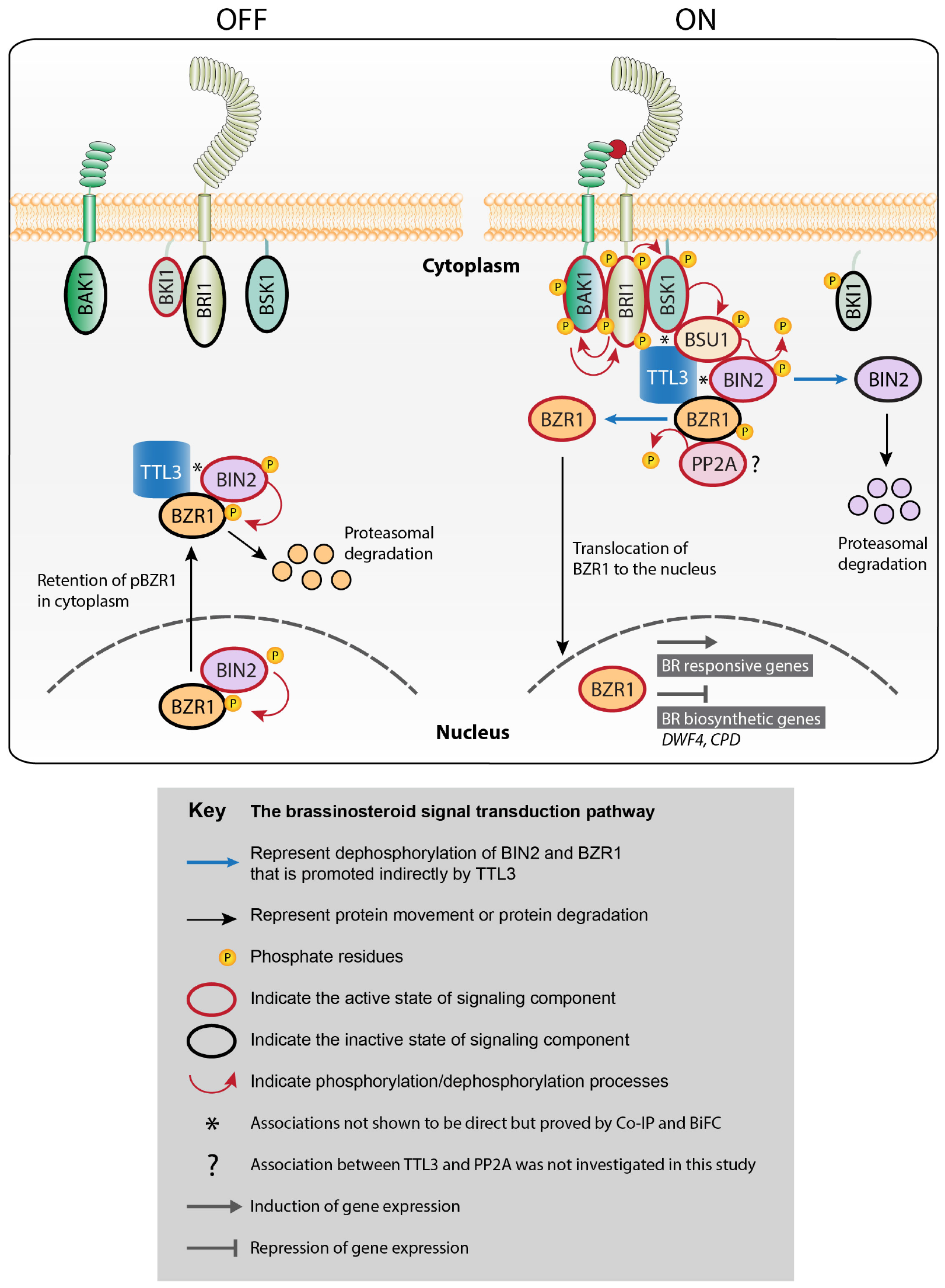
A Proposed model to illustrate how TTL3 mediates a scaffolding mechanism to optimize brassinosteroid signaling. The present study reveals that TTL3 acts as a positive regulator of brassinosteroid (BR) signaling. Our data show that TTL3 presents mainly a cytoplasmic localization in the absence of BR but accumulates at the plasma membrane in response to BR perception. We show that TTL3 directly interacts with BRI1, BSU1 and BZR1, and associates with BSK1 and BIN2 to assemble a BR perception protein complex at the plasma membrane in order to optimize the BZR1 dephosphorylation and active BR signaling. Inactive pathway (OFF) represents the absence of BR and activated pathway (ON) the presence of BR. OFF: In the absence of BR, BRI1 is inactivated by BKI1 and the other plasma membrane components, BAK1 and BSK1, do not associate with BRI1 to form an active complex. In the cytoplasm and in the nucleus, BIN2 phosphorylates BZR1, promoting the inhibition of its DNA-binding activity and its cytoplasmic retention and subsequent degradation in a proteasome-dependent manner. ON: BR binding to the extracellular domain of BRI1 induces not only its dissociation with BKI1 but also its association with the co-receptor BAK1, which functions as a co-receptor of BR. This leads to the activation of BRI1 by trans-phosphorylation events. BAK1 activated BRI1 phosphorylates BSK1 kinase and also causes re-localization of TTL3 to the plasma membrane. There, TTL3 preferentially associates with the active (phosphorylated) BSU, facilitating the dephosphorylation and inactivation of BIN2, which is subsequently degraded by the proteasome. This BIN2 inactivation causes BZR1 dephosphorylation by PP2A and translocation to the nucleus to regulate the transcription of BR target genes. TTL3, TETRATRICOPEPTIDE THIOREDOXIN-LIKE 3; BRI1, BRASSINOSTEROID INSENSITIVE 1; BAK1, BRI1-ASSOCIATED KINASE 1; BKI1, BRI1 KINASE INHIBITOR 1; BSK, BRI1 SUBSTRATE KINASE; BSU1, BRI1 SUPPRESSOR 1; BIN2, BRASSINOSTEROID INSENSITIVE 2; PP2A, PROTEIN PHOSPHATASE 2A; BZR1, BRASSINAZOLE-RESISTANT 1.

Although in current models of BR signaling phosphorylation/de-phosphorylation of transcription factors take place exclusively in the cytoplasm and the nucleus ^559^, a survey of the literature provides evidence that the plasma membrane could be an active site of BR signaling, from perception of the hormone to dephosphorylation of the transcription factors: (1) a significant amount of phosphorylated BZR1 located at the plasma membrane is greatly reduced upon BR treatment ^48^; (2) several BSKs that are plasma membrane-bound interact with BIN2, suggesting that dephosphorylation of BZR1 and BIN2 is also taking place at the plasma membrane ^46,54^; (3) BSK1 has been identified as an interactor of BZR1 using non-targeted proteomics, which led the authors to propose that BR-signaling components exist in the plasma membrane as a multi-protein complex ^55^. We show that BZR1 and BSK1 weakly interact at the plasma membrane and that coexpression of TTL3 greatly increases this association, supporting a role of the plasma membrane in BR signaling.

BR signaling mediated by TTL3 resembles that of Wnt/β catenin signaling which controls many biological processes in metazoans, including cell fate determination, cell proliferation, and stem cell maintenance ^60–62^. In both cases, extracellular ligands are perceived by transmembrane receptors and the signal is transduced through phosphorylation events where GSK3 type kinases phosphorylate effector proteins (either β-catenin in Wnt/β catenin signaling or BZR1/BES1 in BR signaling), resulting in their stabilization or degradation ^5,60–62^. Interestingly, an essential component of this so-called destruction complex involves the central scaffold protein Axin1, which, similar to TTL3, interacts with the core signaling components. In resting conditions GSK3 phosphorylates and degrades β-catenin, although upon Wnt perception the Axin complex is relocalized from the cytoplasm to the plasma membrane, where it suppresses ubiquitination of β-catenin, leading to saturation of complex by accumulation of phospho-β-catenin. As a result, newly synthesized β-catenin can accumulate in the cytosol and translocate to the nucleus, where it promotes transcription ^61,62^.

The basic function of scaffolding proteins is the assembly of signaling components to enhance the efficiency of the signaling cascade by increasing their local concentrations as well as the localization of the signaling reaction to a specific area of the cell. This could be particularly important in BR signal components because some of these proteins are expressed at vanishingly low levels like BSU1 and BIN2 ^63,64^ This scaffolding function of TTL proteins might also have a role in enhancing signaling specificity by preventing spurious interactions by BR signaling components and generating BR specificity. This is important because some components of the BR pathway have been reported to participate in signaling pathways different from the BR one. For example, BAK1 and related SERK co-receptors are involved in numerous signaling pathways ^25^ in addition to its role in BR signaling, and BIN2 shows multiple targets that result in different signaling outcomes ^65,66^. Another example is BSK1, which was originally identified as a BR signaling component by proteomic studies ^67^ but was later found regulate also immune signaling ^68^. Because TTL1, TTL3, and TTL4 genes were previously reported to play a role in abiotic stress tolerance and there is an increasing evidence for the co-ordination of BR-promoted growth and abiotic stress responses ^7–9^, we cannot exclude that function of TTL3 (and probably other TTLs) as a scaffold of BR signaling components contribute to this cross-talk.

Our work uncovers an essential component of BR signaling and fills a gap in our understanding of signaling cascades from the plasma membrane to the nucleus in plant cells. The characterization of other possible scaffold proteins, with a function equivalent to TTLs, will be key to understand how signaling components are assembled in other signaling cascades to ensure the timely signal transduction upon perception of extracellular signals.

---

**Supplementary Figure 1.**
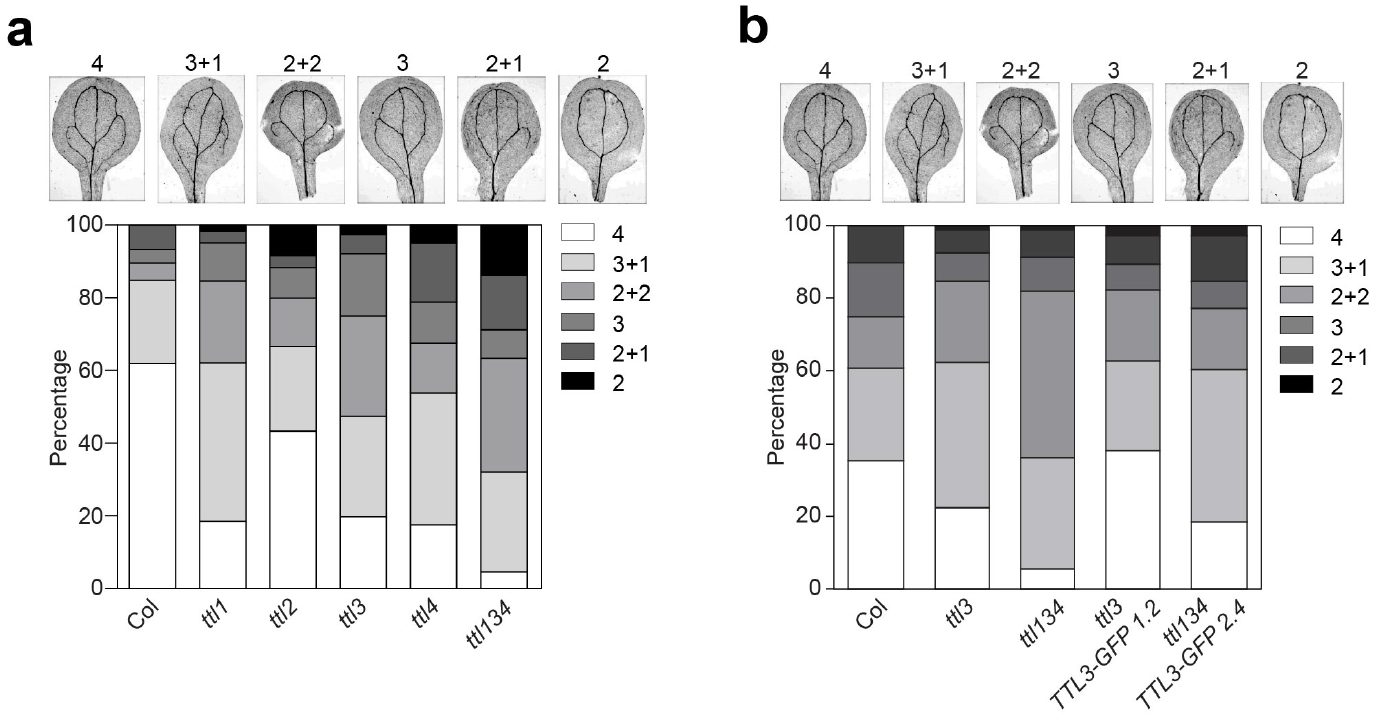
*ttl1, ttl3, ttl4* and *ttl134* are impaired in cotyledon veins pattern formation. **a** *TTL* genes are required for cotyledon vein pattern formation. In wild-type Arabidopsis cotyledons, two types of veins are present: a midvein (or primary vein) and secondary veins that branch from the midvein to form four loops. Increased defects occur if number of loops is reduced when secondary veins do not connect to the base of the midvein or are absent. In *ttl1, ttl3* and *ttl4* mutants, the percentage of seedlings presenting closed loops is clearly reduced. These defects were markedly enhanced in the triple *ttl134* mutant. Categories of cotyledon (embryonic leaves) vein patterns analyzed by stereomicroscope in two-week-old seedlings (upper panel). Cotyledons from two-weeks-old seedlings were observed under a light microscope to identify vascular patterning and the percentage of cotyledons displaying each venation pattern category is depicted at the right side of the panel (bottom panel) were quantified. Approximately 200 cotyledons were analyzed per genotype. **b** *TTL3p::TTL3g-GFP* expression complements cotyledon vein pattern phenotypes of *ttl3* and *ttl134* mutants. Cotyledons vein patterns were analyzed as described in (**a**).

**Supplementary Figure 2.**
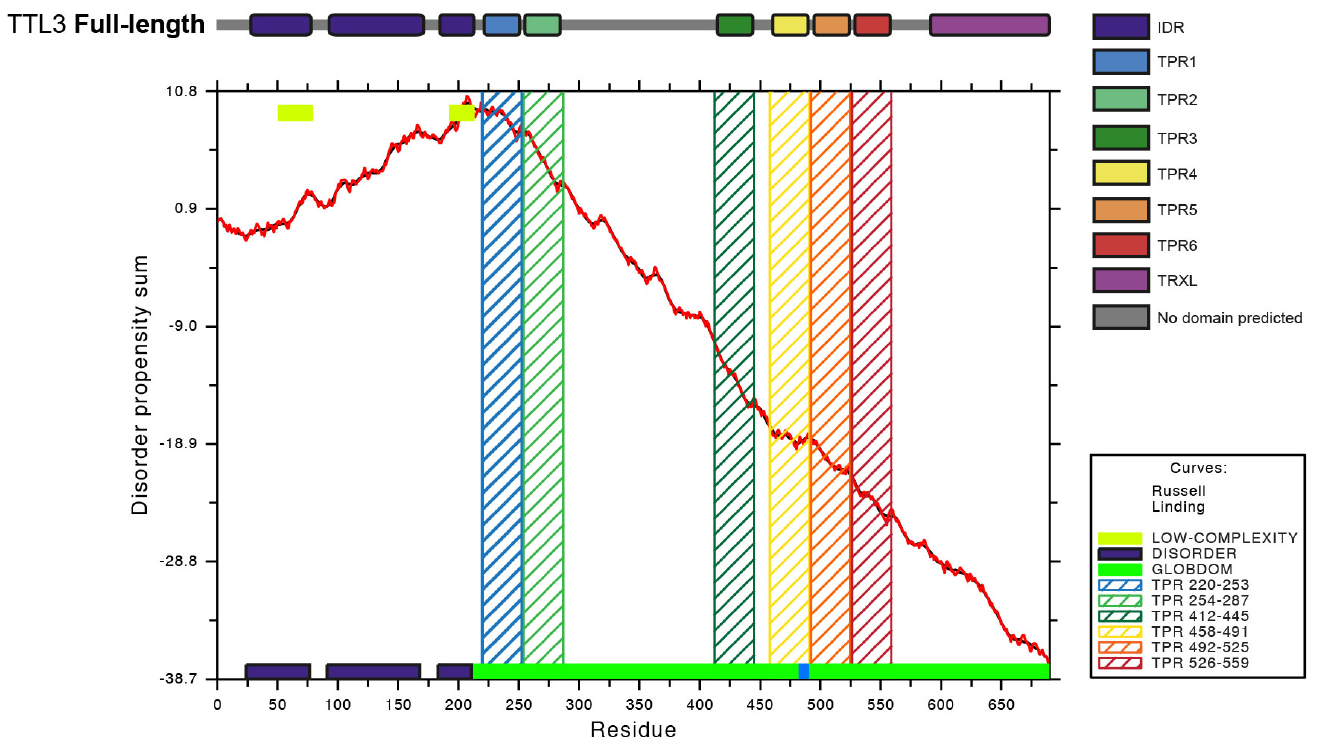
TTL3 presents an intrinsically disorder region (IDR) at the N-terminus. The first 200 amino acids of TTL3 present a disordered structure. Schematic representation of full-length TTL3 protein (upper panel) as described in **Figure 1c** and graphical representation of the TTL3 structure using GlobPlot 2, available in the web page (http://globplot.embl.de/). Disorder propensity sum: up-hill regions correspond to predicted protein disorder (shown in blue) and down-hill regions correspond to putative domains (shown in green). SMART/Pfam domains are also shown. **C.** Schematic representations of the full-length and the truncated version of TTL3 without the first 203 amino acids used for expressing the protein in yeast.

**Supplementary Figure 3.**
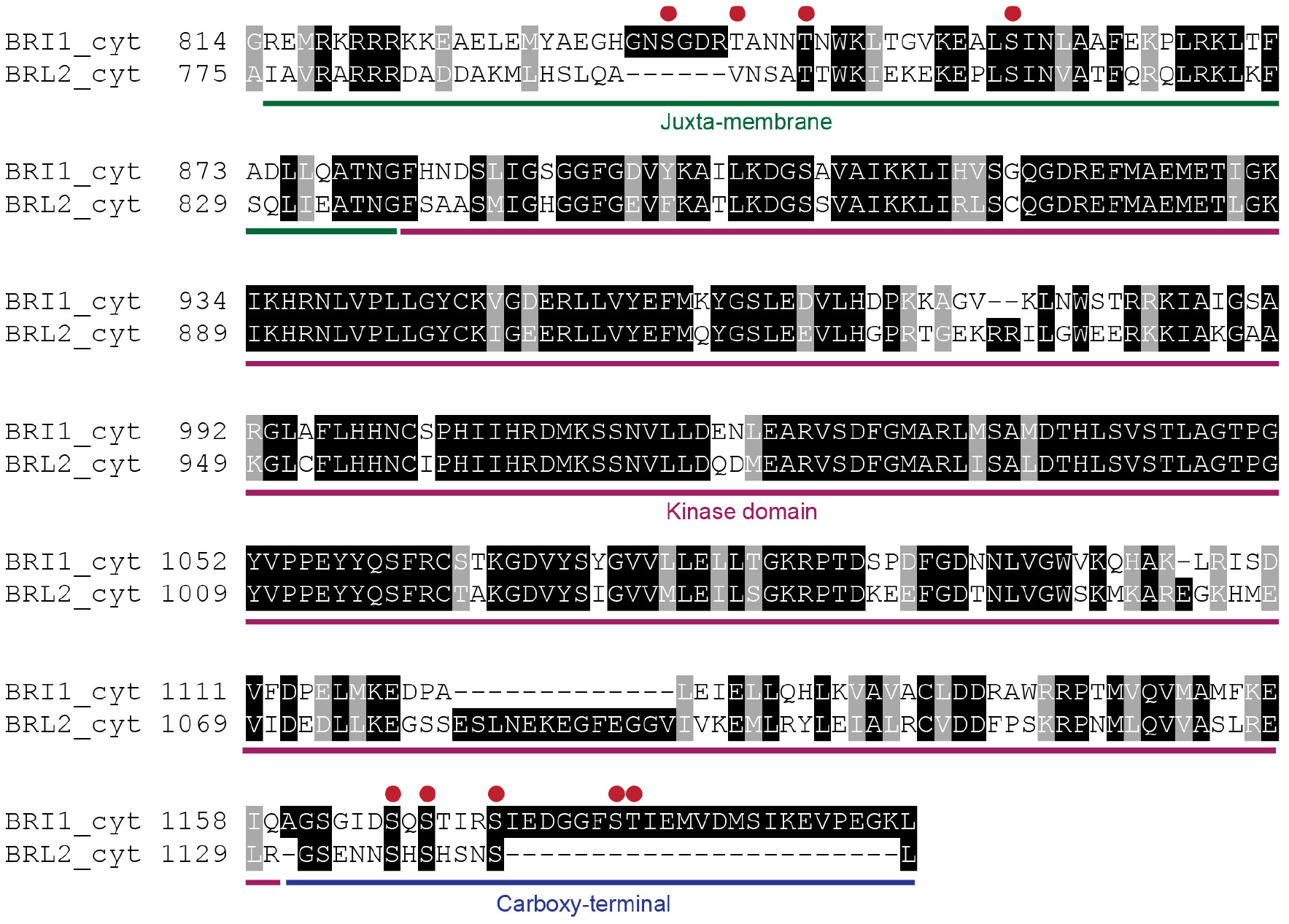
Protein sequence alignment of BRI1 and BRL2 cytoplasmic domain. The protein sequences *Arabidopsis thaliana* BRI1 (*AT4G39400*) and BRL2 (AT2G01950) were retrieved from the TAIR database. The multiple sequence alignment of the cytoplasmic domain protein sequences of BRI1 (residues 814-1196) and BRL2 (residues 775-1143) was performed using T-Coffee alignment package (http://tcoffee.crg.cat/apps/tcoffee/do:regular) and formatted using the Boxshade tool (http://www.ch.embnet.org/software/BOXform.html). The juxta-membrane, the kinase and the carboxyl-terminal domains, are underlined in green, magenta and blue, respectively. The Serine/Threonine residues of the juxta-membrane and carboxyl-terminal domains that were substituted to Aspartic Acid in the BRI1cyt^JMCT9D^ protein are indicated by red dots. Black and gray boxes highlight identical and similar amino acids, respectively.

**Supplementary Figure 4.**
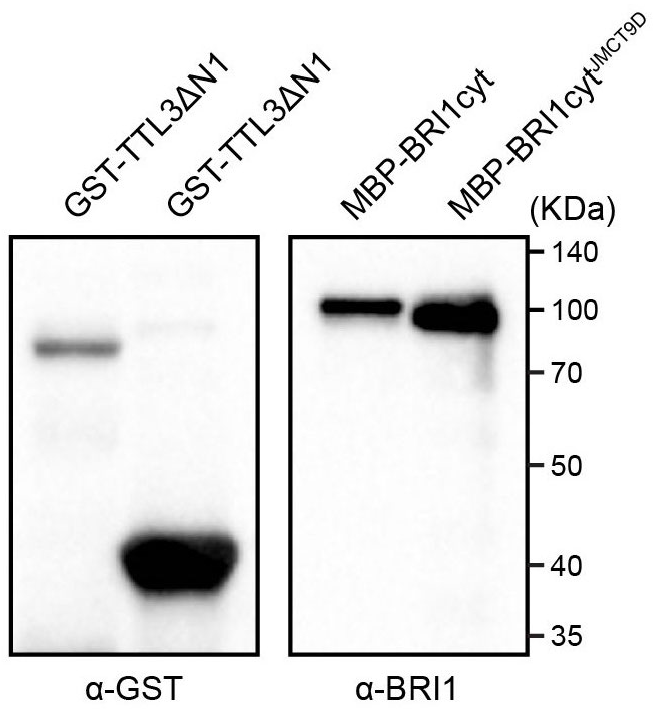
Purified GST-TTL3ΔN1, GST-TTL3ΔN3, MBP-BRIIcyt and MBP-BRI1cyt^JMCT9D^ used for the GST Pull-down assays described in Figure 1d. GST and MBP tagged proteins were expressed in *E. coli*, purified and analyzed by western blot. GST-TTL3ΔN1 and GST-TTL3ΔN3 were detected with anti-GST antibody. MBP-BRI1cyt and MBP-BRI1cyt^JMCT9D^ were detected using specific anti-BRI1 antibodies ^71^.

**Supplementary Figure 5.**
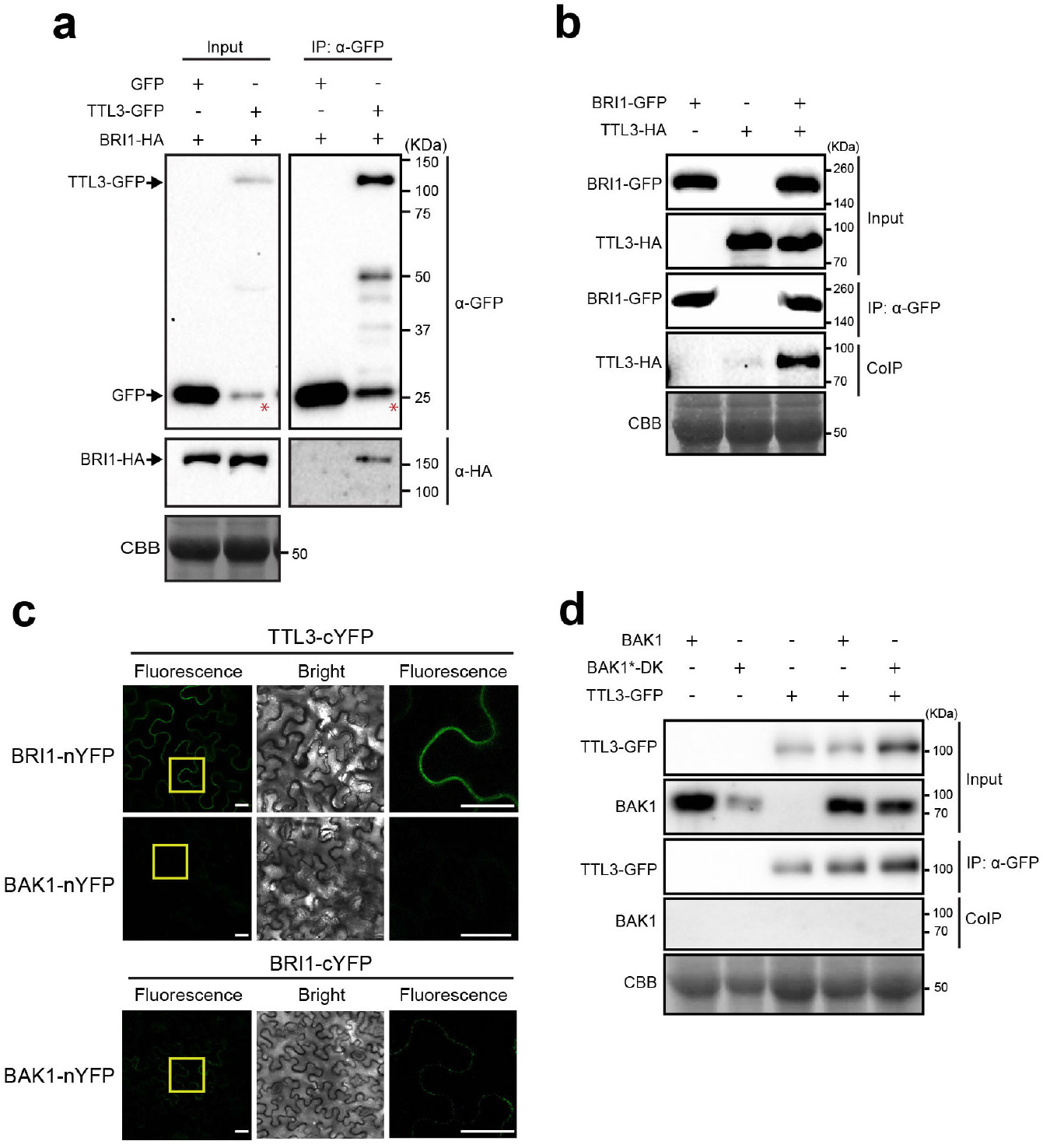
TTL3 specifically associates with BRI1 but not with BAK1 or free GFP. **a** BRI1-HA co-immunoprecipitates with TTL3-GFP but not with free GFP. Epitope Tagged proteins were transiently expressed in *N. benthamiana*, immunoprecipitated and TT3-GFP tagged protein and free GFP was immunoprecipitated using anti-GFP Trap beads. Total (input), immunoprecipitated (IP) and Co-Immunoprecipitated (CoIP) proteins were analyzed by western blotting. Equal loading was confirmed by Coomassie blue staining (CBB) of input samples. Free GFP was used as negative control for Co-IP. TTL3-GFP and free GFP were detected with anti-GFP antibody and BRI1-HA was detected with anti-HA antibody. Asterisks indicate GFP that results from proteolytic cleavage of TTL3-GFP. **b** TTL3-HA co-immunoprecipitates with BRI1-GFP. Epitope Tagged proteins were transiently expressed in *N. benthamiana*, immunoprecipitated and analyzed as indicated in **a**. BRI1-GFP and TTL3-HA were detected with anti-GFP and anti-HA antibodies, respectively. **c** Reciprocal BiFC experiments confirm the association of TTL3 with with BRI1 but not with BAK1. Leaves of *N. benthamiana* were infiltrated with the *Agrobacterium* strains harboring constructs to express TTL3 and BRI1 proteins fused to the C-terminus of the YFP and, BRI1 and BAK1 proteins fused to the N-terminus of the YFP. Using the same settings in the confocal microscope, YFP fluorescence is observed when TTL3-cYFP is co-expressed with BRI1-nYFP, but no YFP fluorescence is detected when TTL3-cYFP is co-expressed with BAK1-nYFP. A weak YFP fluorescence is observed when BRI1-cYFP is co-expressed with BAK1-nYFP. From left to right columns, images show BiFC YFP fluorescence in green, bright field, and 4× magnification of BiFC YFP fluorescence of the region delimited by the yellow square. Scale bars represent 20 μm. All experiments were repeated at least three times with similar results. **d** TTL3 does not co-immunoprecipitate BAK1 or BAK1*-DK (dead kinase D416N). BAK1, BAK1*-DK (dead kinase D416N) and TTL3-GFP were transiently expressed in *N. benthamiana.* Samples were immunoprecipitated and analyzed as indicated in **a**. TTL3-GFP and BAK1 were detected with anti-GFP and anti-BAK1 antibodies, respectively.

**Supplementary Figure 6.**
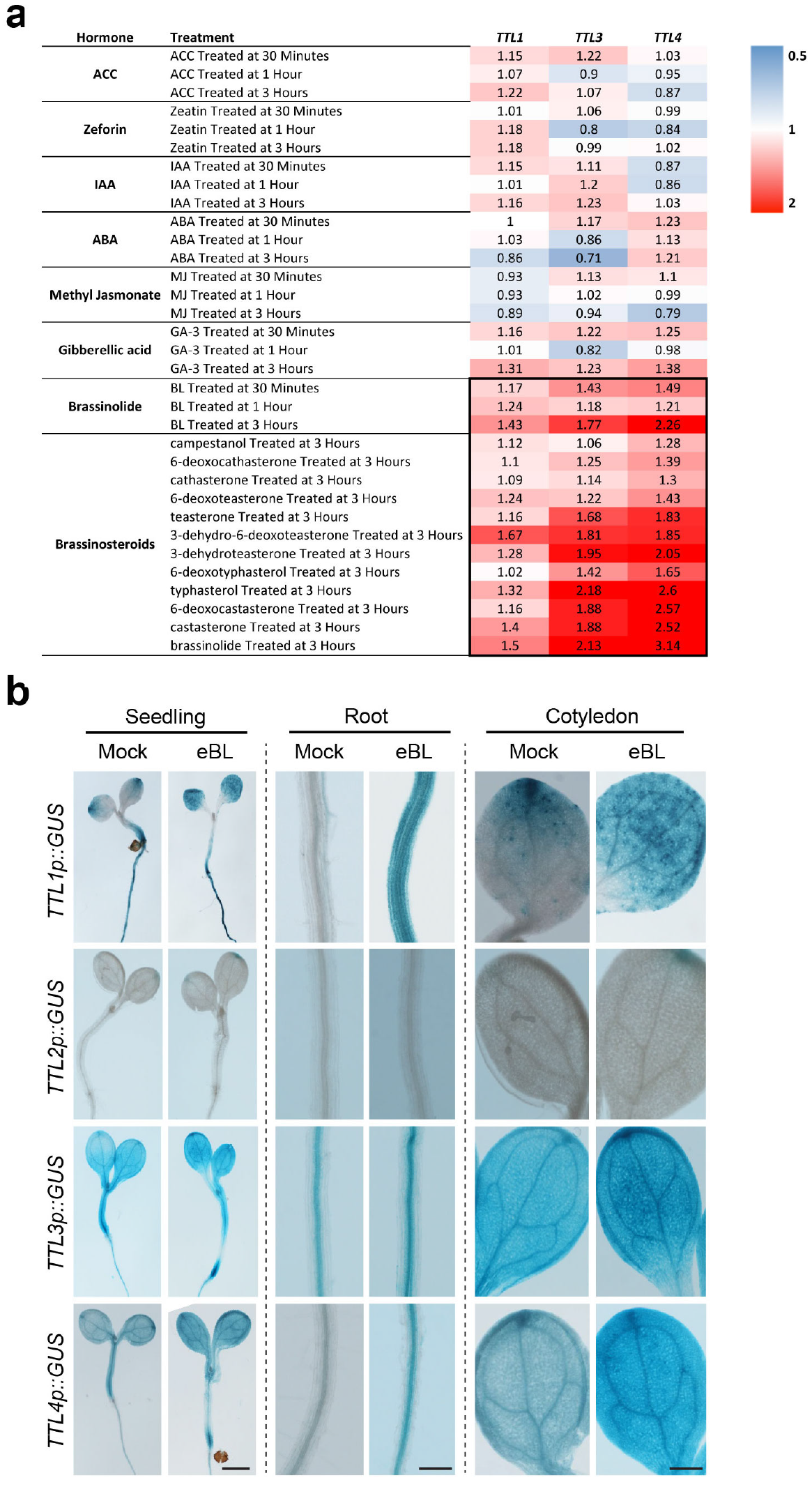
The expression of *TTL1, TTL3*, and *TTL4* are specifically induced by BRs. **a** Heatmap representing the expression responses to hormone treatment of *TTL1, TTL3* and *TTL4. TTL2* was not included due to its low expression in vegetative organs. Expression levels *of TTL* genes are represented as the fold-change relative to the mock treatment. Red colors represent gene inducted and blue colors represent gene repressed in response to the indicated hormone. Gene expression data was retrieved from Arabidopsis eFP Browser (Hormone Series) web site available from the following link: http://bar.utoronto.ca/efp/cgi-bin/efpWeb.cgi (Winter et al. 2007). **b** *TTL1, TTL3*, and *TTL4* promoters are activated by eBL. Histochemical analysis of *TTL* promoters-GUS reporter in control conditions and after exogenous eBL application. Four-day-old seedlings grown in control medium were transferred to a medium containing 0.2 μM eBL for 24 hours and then stained for GUS activity. Scale bars represent 1 mm in seedlings and 200 μm in roots and cotyledon.

**Supplementary Figure 7.**
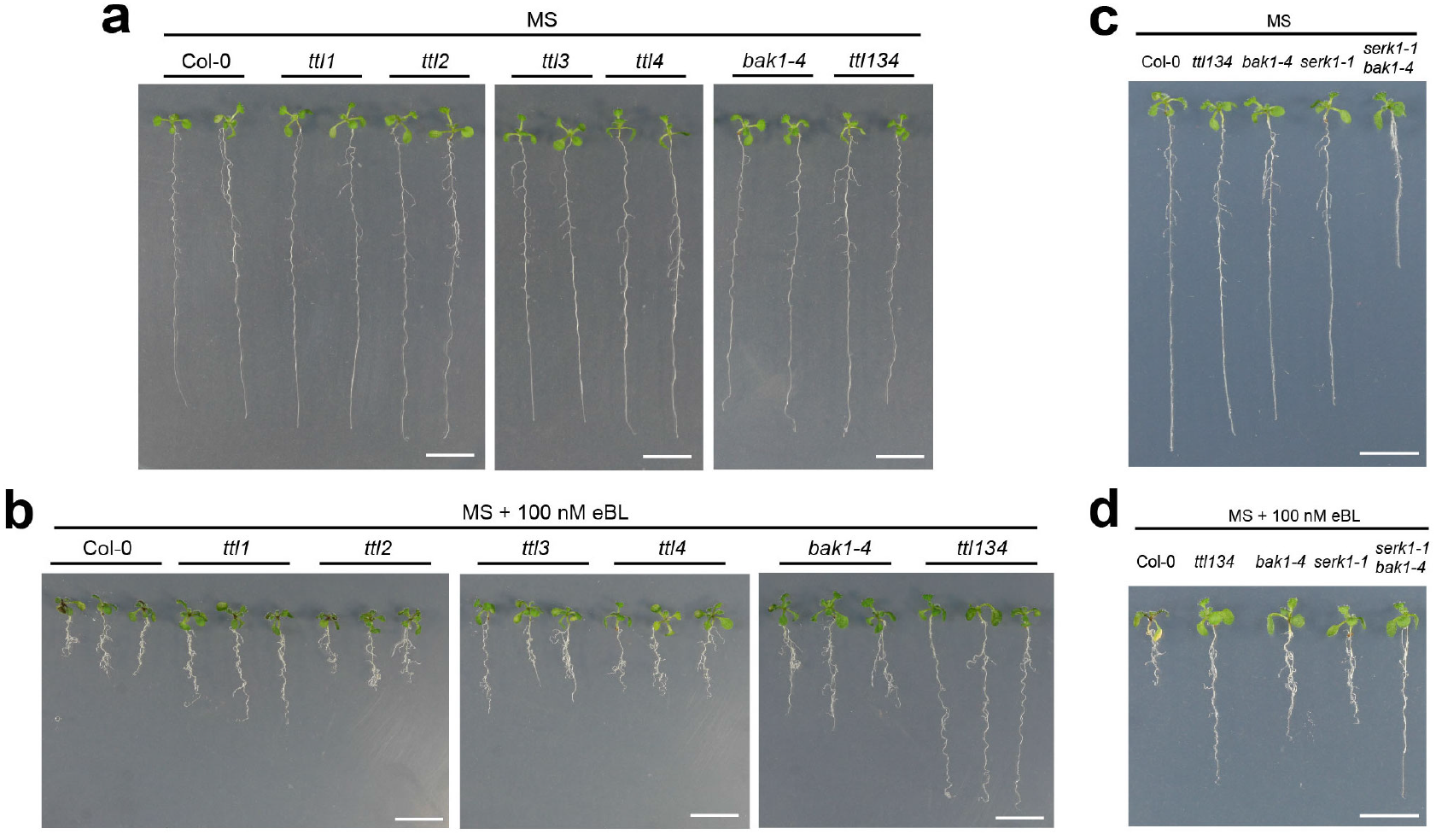
*ttl1, ttl3, ttl4* and *ttl134* show root growth hyposensitivity to BR. **a-b** Root length of WT Col-0, single *ttl* mutants, the triple *ttl134* and the BR perception mutant *bak1-4* in response to eBL. Seedlings were grown in long days for 4 days in half-strength MS agar solidified medium and then transferred to half-strength MS agar solidified medium (**a**, MS) or half-strength MS agar solidified medium supplemented with 100 nM of Brassinolide (**b**, MS + 100 nM eBL) and photographed 6 days later. Scale bars represent 1 cm. **C**. Photographed seedlings are representative of the phenotype observed in the total analyzed replicates, n≥35 seedlings per experiment. The experiment was repeated three times with similar results. **c-d** Root length responses to eBL of wild-type Col-0, *ttl134* and BR perception mutants. Seedlings were grown and root length was analyzed as described in a. Photographed seedlings are representative of the phenotype observed in the total analyzed replicates, n=30 seedlings per experiment. The experiment was repeated three times with similar results.

**Supplementary Figure 8.**
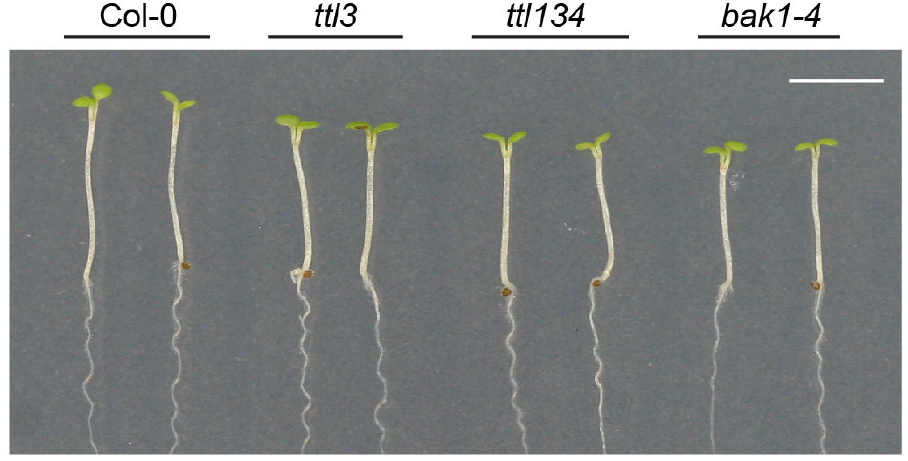
Defective hypocotyl elongation in *ttl* mutants. Col-0, *ttl3, ttl134* and *bak1-4* seedlings were grown for 4 days in long-day photoperiod in half-strength MS agar solidified medium. Seedling with the same size were then placed in the dark and photographed 3 days later. Scale bar represents 5 mm. Photographed seedlings are representative of the phenotype observed in the total analyzed replicates, n = 80 seedlings per experiment. The experiment was repeated twice with similar results.

**Supplementary Figure 9.**
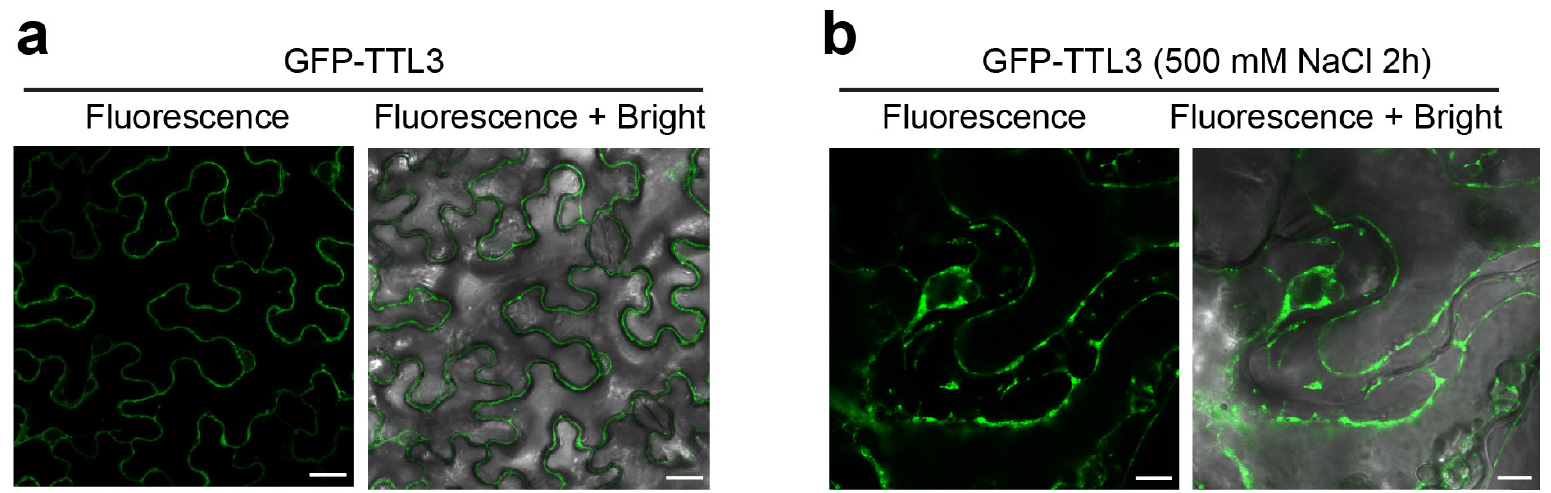
TTL3 presents a cytoplasmic/plasma membrane sub-cellular localization. **a** Confocal microscopy images showing *N. benthamiana* transiently expressing GFP-TTL3 indicate a main cytoplasmic localization. Scale bars represent 20 μm. **b** *N. benthamiana* leaves expressing GFP-TTL3 after plasmolysis indicates plasma membrane localization. Leaf cells were plasmolyzed using 500 mM NaCl for 2 hours. Confocal microscopy images show that GFP signal remains in the retracted Hechtian strands at the plasma membrane bound to the cell wall. Scale bars represent 10 μm.

**Supplementary Figure 10.**
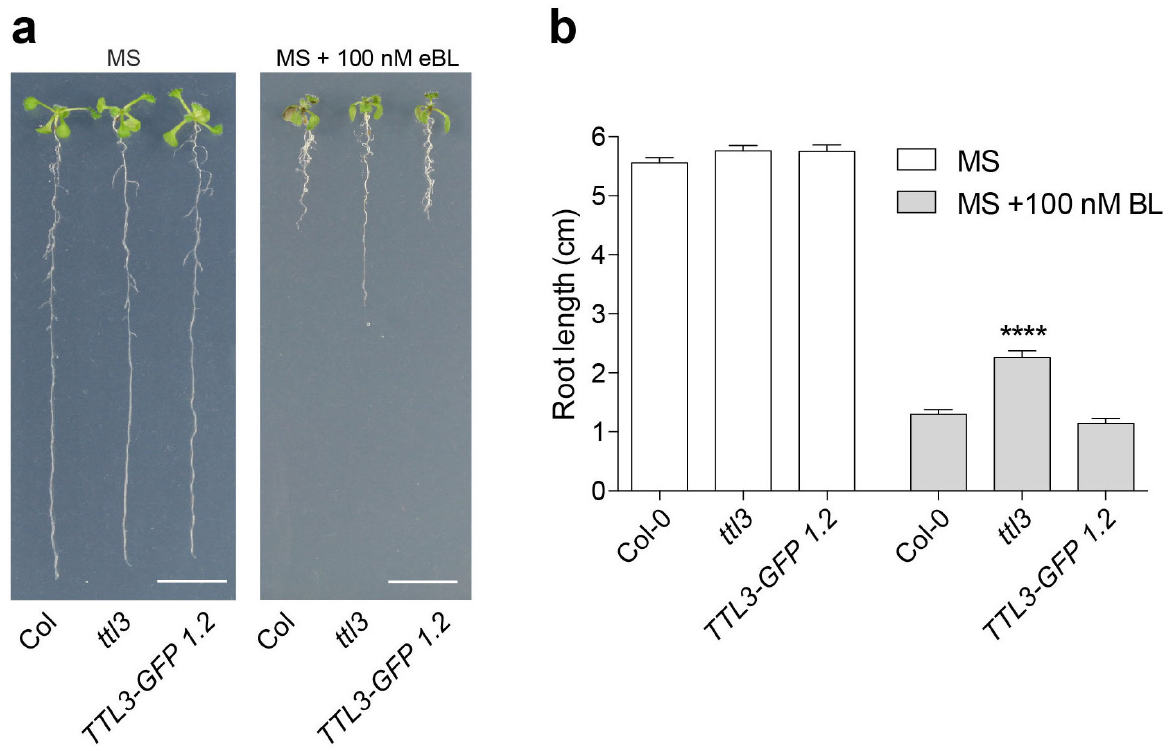
*TTL3-GFP* 1.2 complements the root length phenotype of *ttl3* in response to eBL treatment. **a** Seedlings were grown for 4 days in half-strength MS medium and then transferred to MS medium or half-strength MS medium supplemented with 100 nM of epiBrassinolide (eBL). Seedlings were photographed 6 days later. Scale bar represents 1 cm. **b** Statistical analysis of root length of Col-0, *ttl3* and the complementation line *TTL3-GFP* 1.2 described in **a**. Asterisks indicate statistical differences significance between the indicated genotype vs Col-0 as determined by the unpaired *t-test* (**** P ≤ 0.0001). Data represent mean values, error bars are SEM, n=30 seedlings per experiment. The experiment was repeated three times with similar results.

**Supplementary Figure 11.**
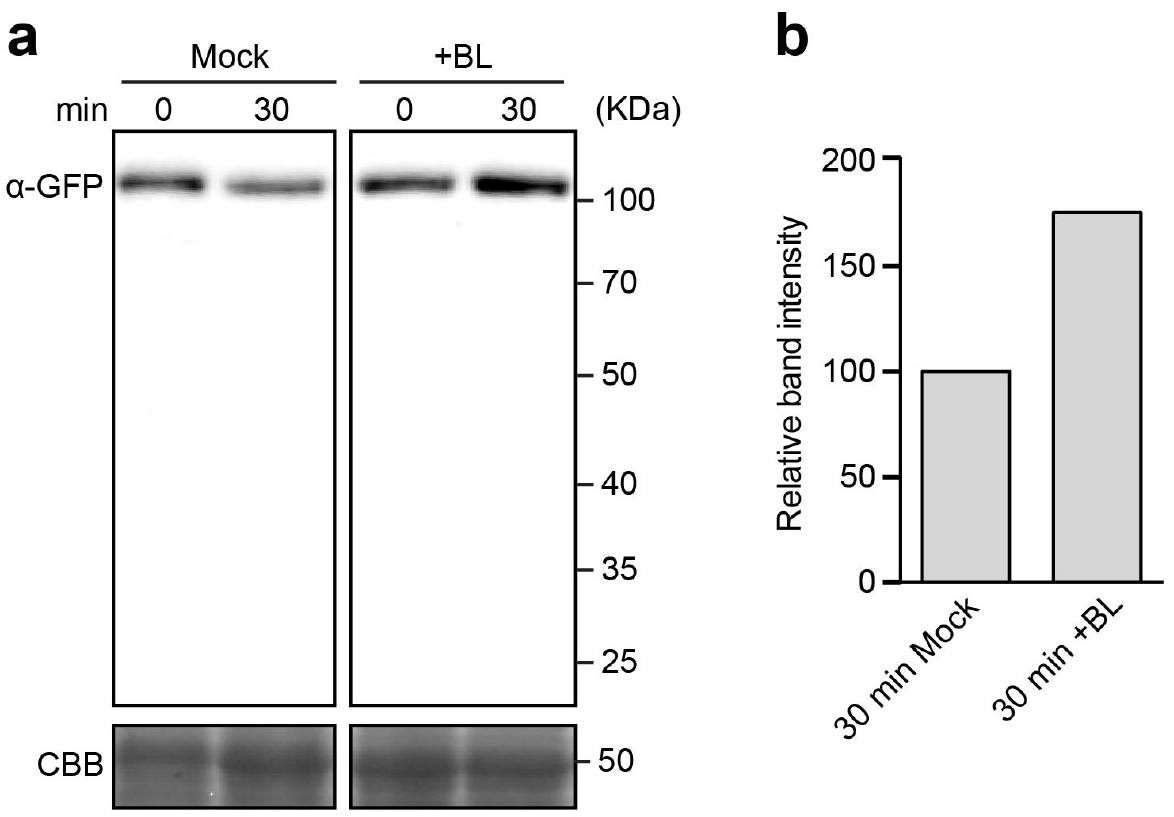
Western blot analyses revels that eBL treatment induces TTL3-GFP protein stabilization, and that there in no degradation-products of TTL3-GFP in Arabidopsis *TTL3-GFP* 2.4 line. **a** Western blot analysis of TTL3-GFP protein in 3-day-old Arabidopsis seedlings of *TTL3-GFP* 2.4. Full scan data of the immunoblot is shown demonstrate that there is no degradation-products of TTL3-GFP in the Arabidopsis *TTL3-GFP* 2.4 line. Seedlings of *TTL3-GFP* 2.4 line were grown in control conditions (mock) or treated with 1 μM eBL for 30 minutes. **b** Graphical representation of the normalized TTL3-GFP +BL protein levels expressed as relative abundance to the amount of the TTL3-GFP mock (arbitrarily set at 100). Intensities of the TTL3-GFP protein bands (**a** top panel) and the Coomassie blue-stained gel (**a** bottom panel) were quantified using ImageJ software (http://rsb.info.nih.gov/ij). Image shows the results from one representative experiment. Four independent experiments were performed with similar results.

**Supplementary Figure 12.**
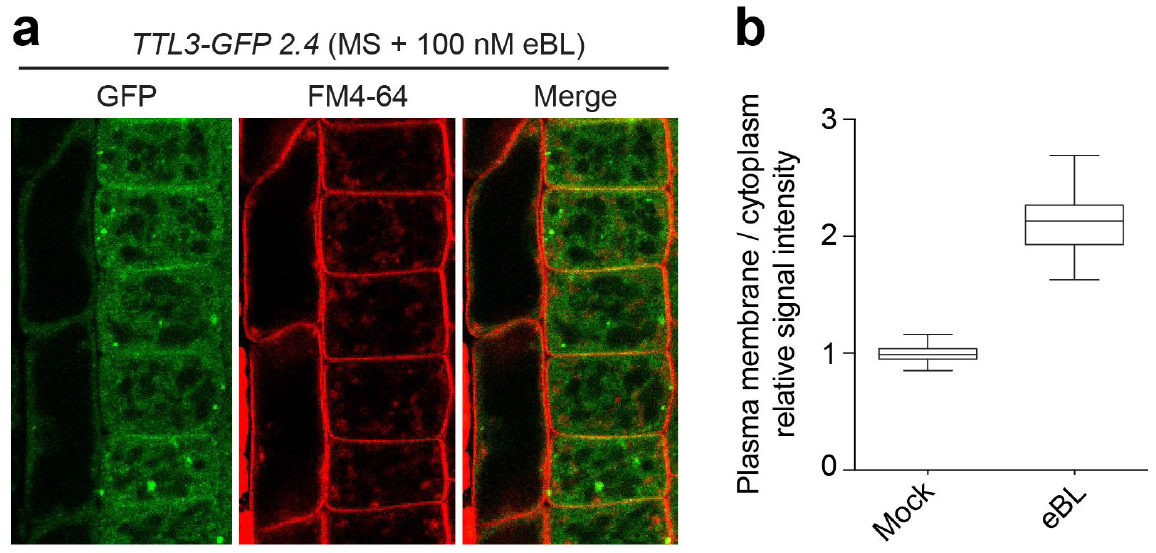
BRs regulate the cytoplasmic/plasma membrane localization of TTL3. **a** Confocal microscopy image showing the localization of TTL3-GFP fluorescence in epidermal cells from root meristematic zone of 4-day-old *TTL3-* GFP 2.4 after 1 hour of 1 μM eBL treatment. Red channel shows the plasma membrane stained with FM4-64. Scale bar represents 10 μm. **b** Quantification of the GFP signal in plasma membrane vs cytoplasm. To measure the ratio between plasma membrane and cytoplasmic signals, a small area of fixed size (8 pixels) was drawn, and measurements of integrated densities were taken from representative areas within the plasma membrane and cytoplasm of each cell. To delimitate the plasma membrane area, FM4-64 was used to stain the cells as depicted in a. Average ratios between plasma membrane and cytoplasmic signal intensities were calculated based on measurements from 3 cells per plant. 10 plants analyzed. N=30. This experiment was repeated twice with similar results.

**Supplementary Figure 13.**
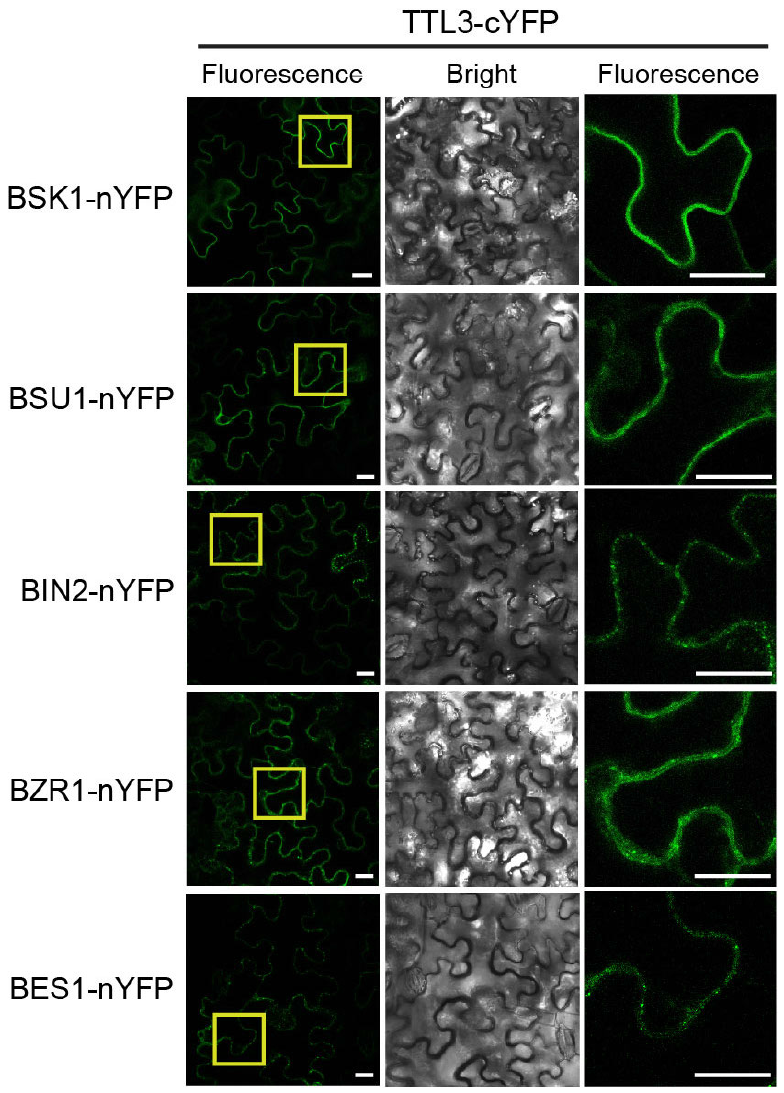
TTL3 associates with BSK1, BSU1, BIN2, BZR1 and BES1 by BiFC Reciprocal BiFC experiments. Reciprocal BiFC experiments confirm the interaction of TTL3 with BSK1, BSU1, BIN2, BZR1 and BES1. Leaves were co-agroinfiltrated with the *Agrobacterium* strain harboring a construct to express the TTL3 protein fused to the C-terminus of the YFP, and the BSK1, BSU1, BIN2, BZR1 and BES1 proteins fused to the N-terminus of the YFP. By confocal microscopy, YFP fluorescence is observed when TTL3-cYFP is co-expressed with BSK1-nYFP, BSU1-nYFP, BIN2-nYFP, BZR1-nYFP and BES1-nYFP. From left to right columns, images show BiFC YFP fluorescence in green, bright field, and 4X magnification of BiFC YFP fluorescence of region delimited by the yellow square. Scale bars represent 20 μm.

**Supplementary Figure 14.**
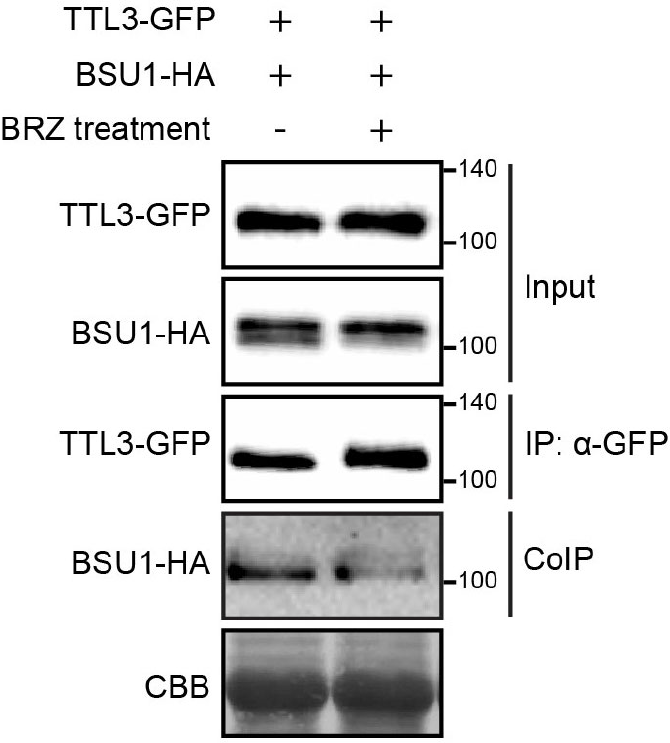
TTL3 preferentially associates with active BSU1 by CoIP. TTL3 preferentially associates with active BSU1. BSU1-HA and TTL3-GFP proteins were transiently expressed in *N. benthamiana* pre-treated with mock solution or with 5 μM BRZ for 48h. TTL3-GFP was immunoprecipitated with anti-GFP Trap beads. Total (input), immunoprecipitated (IP) and Co-Immunoprecipitated (CoIP) proteins were analyzed by western blotting. Equal loading was confirmed by Coomassie blue staining (CBB) of input samples. GFP-TTL3 and BSU1-HA were detected with anti-GFP and anti-HA antibody, respectively. The Co-IP shows an enrichment of the lower BSU1 band, despite the decrease in the input caused by BRZ.

**Supplementary Figure 15.**
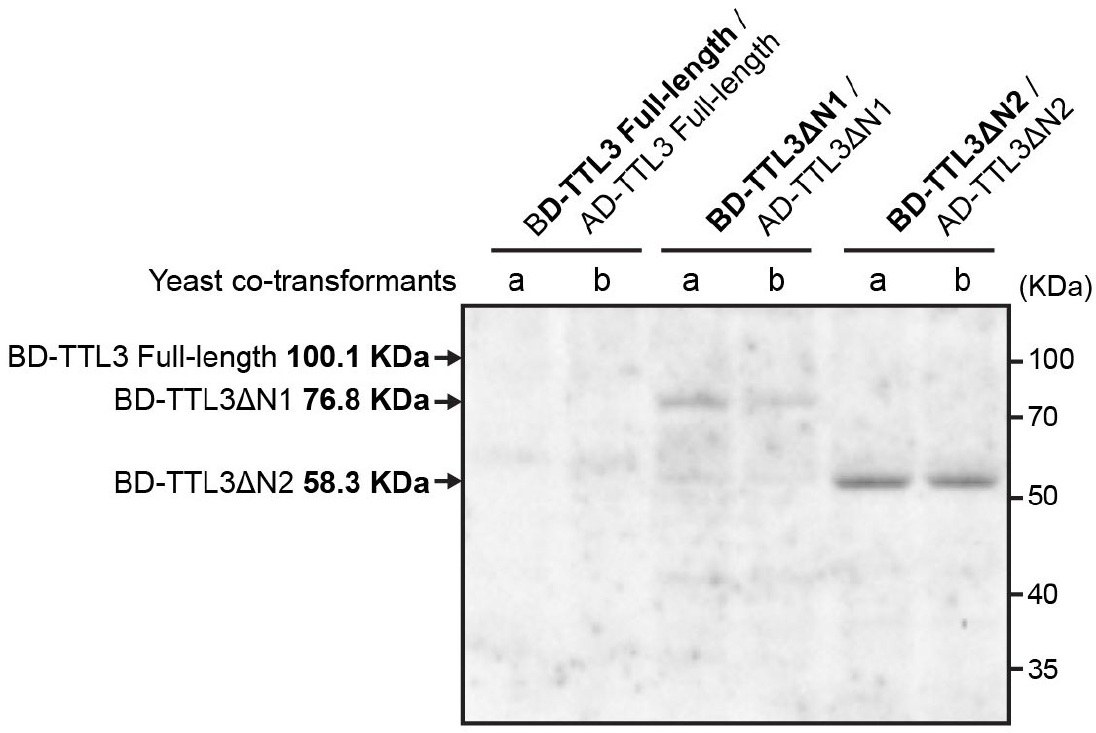
TTL3 N-terminus negatively affects the stabilization of TTL3 protein in yeast heterologous system. Protein extracts of two independent yeast co-transformants (a and b) for each bait-prey plasmid combination were resolved in polyacrylamide/SDS-Page gels and analyzed by western blot using a anti-Myc Tag monoclonal antibody. Myc Tag is transcriptionally fused to BD-fused protein (in bold). The expected molecular size of BD-TTL3 Full-length, BD-TTL3ΔN1 and TTL3ΔN2 is represented in the figure.

**Supplementary Figure 16.**
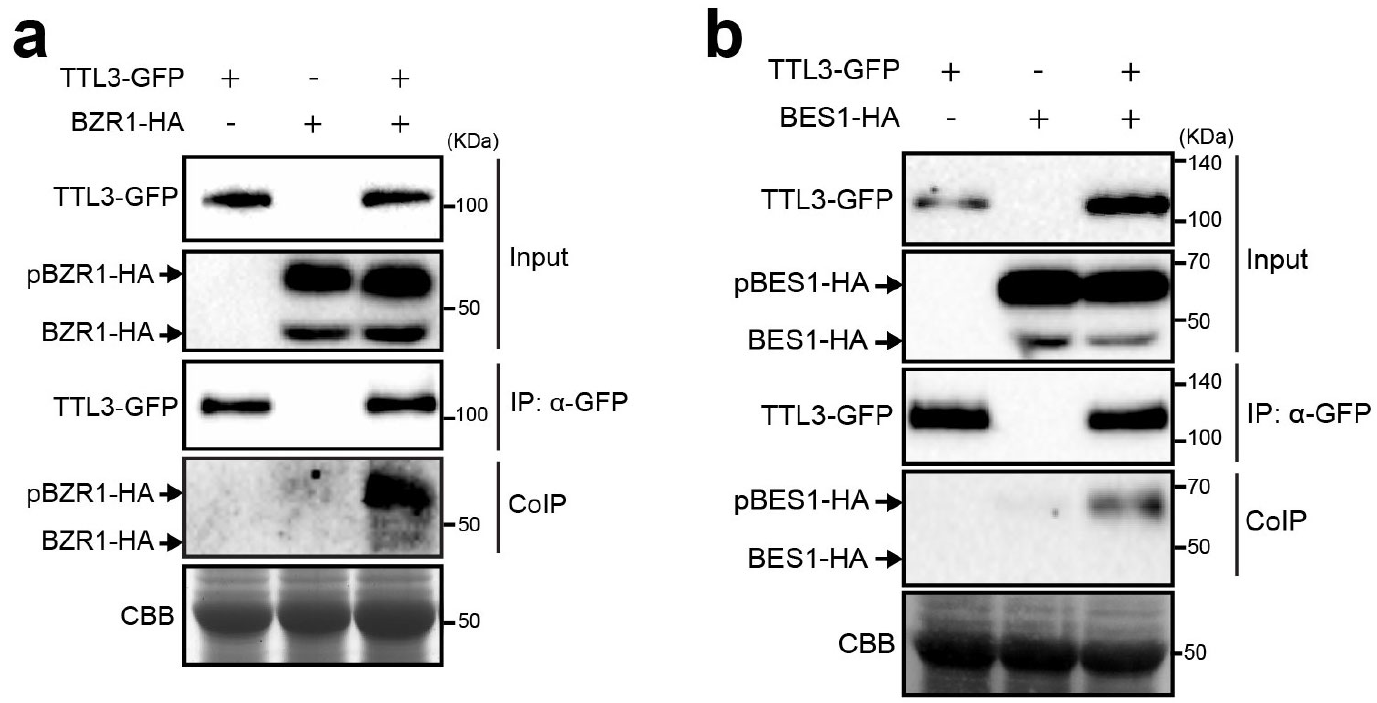
TTL3 preferentially associates with phosphorylated (inactive) form of BZR1 and BES1 by CoIP. **a** Co-immunoprecipitation of BZR1 with TTL3 indicates a preferential association of TTL3 with pBZR1. BZR1-HA and TTL3-GFP were transiently expressed in *N. benthamiana* and TTL3-GFP was immunoprecipitated with anti-GFP Trap beads. Total (input), immunoprecipitated (IP) and Co-Immunoprecipitated (CoIP) proteins were analyzed by western blotting. Equal loading was confirmed by Coomassie blue staining (CBB) of input samples. TTL3-GFP and BZR1-HA were detected with anti-GFP and anti-HA, respectively. The upper band corresponds to phosphorylated BZR1 (pBZR1-GFP) and the lower one to dephosphorylated BZR1 (BZR1-GFP). **b** BES1 co-immunoprecipitates with TTL3. BES1-HA and TTL3-GFP were transiently expressed in *N. benthamiana*, immunoprecipitated and analyzed by western blot as indicated in a. TTL3-GFP and BES1-HA were detected with anti-GFP and anti-HA, respectively.

## Supplemental Tables

**Table S1.**
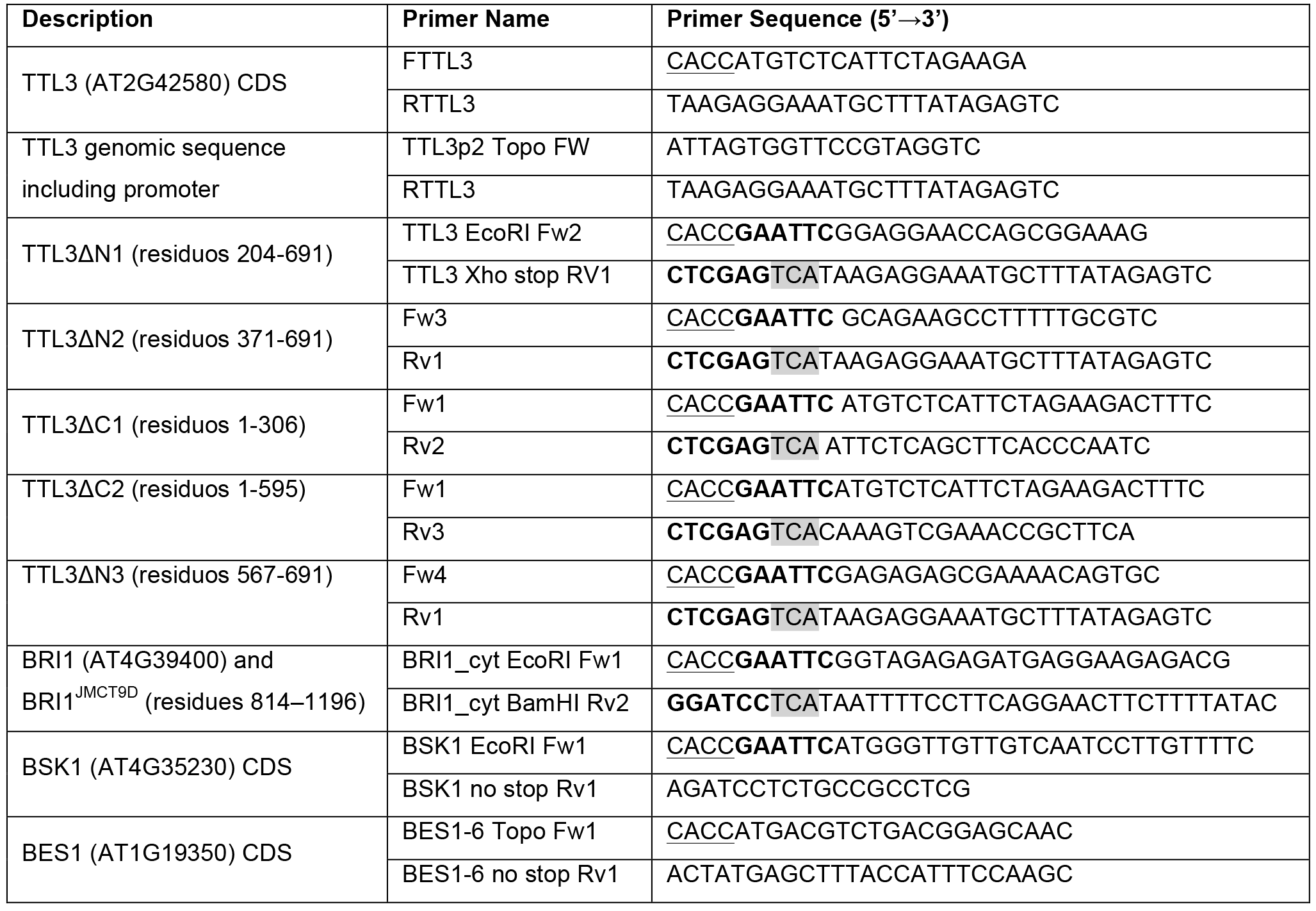
List and description of primers used for cloning into pENTR. Restriction sites included in some primers are highlighted in bold and enzyme is indicated in the primer name. CACC sequence include in Fw primers to clone in pENTR/D-TOPO (Invitrogen) is underlined. STOP codon in primers sequence is highlighted in gray.

**Table S2.**
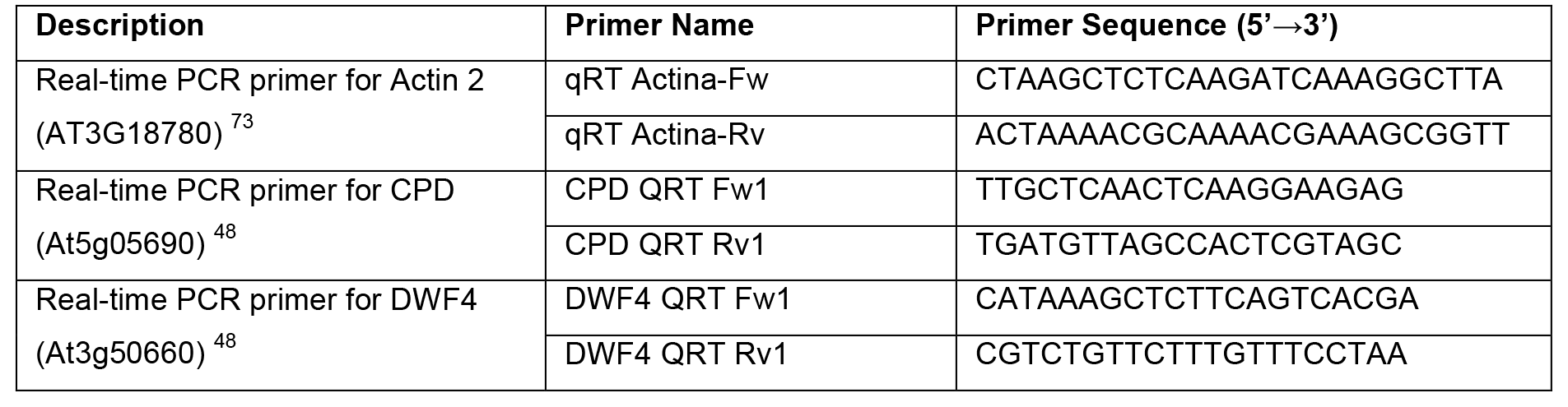
List of primers used for quantitative RT-PCR.

## METHODS

### Plant Material and Growth Conditions

All *Arabidopsis thaliana* plants used in the present study were Columbia-0 ecotype (Col-0). Arabidopsis mutants lines used in this study have been previously described: *ttl1* (AT1G53300) Salk_063943; *ttl2* (AT3G14950) Salk_106516; *ttl3* (AT2G42580) Sail_193_B05; *ttl4* (AT3G58620) Salk_026396; *ttl134: ttl1 ttl3 ttl4* triple mutant ^12^; *bak1-4* (SALK_116202) ^28^; *serk1-1* (SALK_044330) ^74^ and *serk1-1 bak1-4* double mutant (obtained by crossing *serk1-1* with *bak1-4*). Transgenic lines *TTL1p::GUS; TTL2p::GUS; TTL3p::GUS and TTL4p::GUS* ^12^ were also previously described. Generation of transgenic lines *TTL3-GFP 1.2 (TTL3p::TTL3g-GFP* line 1.2 in *ttl3* background) and *TTL3-GFP 2.4 (TTL3p::TTL3g-GFP* line 2.4 in *ttl134* background) is described in the **Generation of Transgenic Plants** section.

### Plant Manipulation and Growth Conditions

Arabidopsis standard handling procedures and conditions were employed to promote seed germination and growth. Seeds were surface sterilized and cold treated for three days at 4°C. Then, seeds were sowed onto half-strength Murashige-Skoog agar solidified medium (0.6% (w/v) agar for horizontal growth and 1% (w/v) for vertical growth) containing 1.5% sucrose, unless otherwise stated. Plates were placed either vertically or horizontally in a culture chamber at 22 ± 1°C, under cool white light (120 μmol photon m^−2^ s^−1^) with a long-day photoperiod (16-h light/8-h dark cycle) unless otherwise stated. When required, seedlings were transferred to soil after seven days of in vitro growth and watered every two days. In soil, plants were grown in a mixture of organic substrate and vermiculite (4:1 v/v) under controlled conditions: 23 ± 1°C °C, 16-h light/8-h dark cycle (~120 μmol photon m^−2^ s^−1^). Freshly harvested seeds were used for all the phenotypic analysis.

### Plasmid Constructs

A genomic fragment spanning the 1.7 kb *TTL3* promoter (*TTL3p*) region upstream of the start codon and the TTL3 genomic region (*TTL3g*) without stop codon was PCR amplified using the primers detailed in **Table S1** and cloned into pCR8 ENTRY vector (Invitrogen).

The coding DNA sequence (CDS) without the stop codon of *TTL3, BSK1* and *BES1 (BES1-S*, the canonical BES1 isoform), as well as the CDS with stop codon of wild-type BRI1 cytoplasmic domain (residues 814-1196), BRI1 cytoplasmic domain JMCT9D (BRI1cyt^JMCT9D^ residues 250-662), and TTL3 truncated version TTL3ΔN1 (residues 204-691), TTL3ΔN2 (residues 371-691), TTL3ΔN3 (residues 567-691), TTL3ΔC1 (residues 1-306) and TTL3ΔC2 (residues 1-595) was PCR amplified using the primers detailed in **Table S1** and cloned into the *pENTR/D-TOPO* vector using the pENTR Directional TOPO cloning kit (Invitrogen). The pUNI51 (Salk Institute) cDNA clone was used as template to PCR amplify *TTL3* CDS without the stop codon. Total RNA from Arabidopsis Col-0 was used to generate cDNA that was then employed as template to PCR amplify BSK1 CDS without the stop codon. The destination vector pGADT7(GW)BES1 ^32^ was a gift from Salomé Prat (CNB-CSIC), and it was used as template to PCR amplify BES1 (*BES1-S*, the canonical BES1 isoform correspond to the BES1-6 transcript). The expression clone pMAC-flag-BRI1-CD-JMCT9D ^70^ was used, as template to PCR amplification of BRI1cyt^JMCT9D^ and it was a gift from Xiaofeng Wang (College of Horticulture Northwest, A&F University, Yangling Shaanxi).

pENTR vectors including CDS without stop codon of *BRI1, BAK1, BIN2, BSU1* and *BZR1*, were obtained by Gateway BP-reaction (Invitrogen) using an expression clone for each gene of interest (containing *attB* sites) and the pDONR/Zeo vector. Expression clones, used as templates for cloning *BRI1, BAK1*, and *BZR1* in pENTR/D-Topo by Gateway BP-reaction, were previously published ^29,75^ The expression clones used to clone *BSU1* ^63^ and *BIN2* ^32^ in pENTR/D-Topo by Gateway BP-reaction, were a gift from Santiago Mora Garcia (Fundación Instituto Leloir and IIBBA) and Salomé Prat (CNB-CSIC), respectively.

All the resulting pENTR clones were verified by diagnostic PCR, restriction analysis and sequencing. These pENTR clones in combination with the appropriate destination vectors (pDEST) were used to create the final Gateway-expression constructs, by LR-reaction (Invitrogen). The pETG-30A and pETG-30A vectors were provided by the European Molecular Biology Laboratory (EMBL) and were used as pDEST to generate GST and MBP N-terminus fusion proteins for GST-pull-down assays. The pGWB4, 5, 6 and 14, from the pGWBs vectors series, were provided by Tsuyoshi Nakagawa ^76^ (Department of Molecular and Functional Genomics, Shimane University), and were used as pDEST for the transient expression in *N. Benthamiana* in the Co-IP and coexpression assays (pGWB5, 6 and 14) or for generating stable transgenic Arabidopsis lines (pGWB4). The pDEST-GW-VYNE and pDEST-GW-VYCE ^77^ were used for BiFC assays. The Gateway destination vector pUC19(35S::GW-GFP) and pBSSK(35S::GW-HA) were used to transfect protoplast for transient expression and Co-IP assays. The pUC19(35S::GW-GFP) was provided by José Alonso (Department of Plant and Microbial Biology, North Carolina State University), and contains pGWB5 cassette between HindIII-SacI restriction sites in pUC19 vector backbone. The pBSSK(35S::GW-HA) was generated in this work by cloning the pGWB14 cassette between HindIII-SacI in the pBSSK vector backbone. The pGADT7(GW) and pGBKT7(GW) destination vectors were provided by Salomé Prat (CNB-CSIC) and used for yeast two-hybrid assay. All the expression clones were verified by diagnostic PCR and restriction analysis.

### Generation of Transgenic Plants

Expression constructs were transformed into *Agrobacterium tumefaciens* strain GVG3101::pMP90 through electroporation and confirmed by diagnostic PCR. The pGWB4 harboring the *TTL3p::TTL3g-GFP* construct, was transformed into Arabidopsis plants by floral dip ^78^ to generate stable transgenic plants. *TTL3p::TTL3g-GFP* was transformed into both the *ttl3* single mutant and the *ttl134* triple mutant. T3 or T4 homozygous transgenic plants were used in this study.

### Phenotypic Analysis

#### Venation pattern phenotype

Cotyledons (embryonic leaves) from two-week-old seedlings were cleared and observed under a light microscope to analyze vascular patterning and the percentage of cotyledons displaying each venation pattern categories is depicted in **Supplementary Fig. 1**. Approximately 200 cotyledons per genotype were analyzed. Representative images of each observed venation pattern categories were acquired using the Nikon AZ100 Multizoom microscope system.

For clearing cotyledons, the two-week-old seedlings were immersed sequentially in 50% ethanol for 1 hour, 99% ethanol overnight, and 50% ethanol for 1 hour, and finally transferred to ddH_2_O. Seedlings were mounted on slides in 50% glycerol and visualized under a light microscope or using the Nikon AZ100 Multizoom microscope system as described above.

#### BL Sensitivity Determined by Root Growth Inhibition

Seedling were grown vertically in long-day photoperiod for 4 or 5 days in half-strength MS agar solidified medium supplemented with 1.5% (w/v) sucrose, and then transferred to half-strength MS agar solidified medium supplemented with 1, 5% (w/v) sucrose containing either mock (eBL solvent as control) or 100 nM eBL (PhytoTechnology Laboratories) and photographed 6 or 8 days later. The eBL (PhytoTechnology Laboratories) was added from a 5 mM stock solution freshly prepared in 80% (v/v) ethanol.

To determine the eBL sensitivity of Col-0 and mutants, the root length of 10 or 13-day-old seedlings grown vertically as described above was measured and the data were analyzed as described in *“Quantification and Statistical Analysis”* section.

#### BL Sensitivity Determined by Phosphorylation status of BES1

Seedling were grown vertically in long-day photoperiod for 7 days in half-strength MS agar solidified medium supplemented with 1.5% (w/v) sucrose, and then transferred to half-strength MS liquid medium supplemented with 1,5% (w/v) sucrose containing 2.5 μM BRZ (TCI Europe) and grown for 3 more days. To determine the eBL sensitivity of Col-0 and *ttl134*, the seedlings were treated with either mock (eBL solvent as control) or 10 nM eBL (PhytoTechnology Laboratories) and frozen in liquid nitrogen 0, 30 and 60 minutes after the treatment. Total protein was extracted as described in “**Extraction of Total Protein from Arabidopsis**” section and analyzed by immunoblotting using an anti-BES1 antibody (dilution 1:500) (Yu et al., 2011) as described in the “**Western Blot Analysis**” section.

#### Hypocotyl elongation in dark

Freshly harvested seeds were surface sterilized and cold treated for three days at 4°C. Then, seeds were sowed individually onto half-strength Murashige-Skoog 1% (p/v) agar solidified medium containing 1.5% sucrose for vertical growth. Seedlings were grown for 4 days in long-day photoperiod, and then placed in dark condition (vertical growth in a culture chamber at 22 ± 1°C). Seedlings were photographed and hypocotyl length was measured 3 days after placing plates in dark conditions.

### Total RNA Extraction and Semi-quantitative RT-PCR Analysis

Ten-day-old seedlings (10 seedlings per biological replicate) grown for five days on half-strength MS agar solidified medium were transferred to half-strength MS liquid medium supplemented with 1% (w/v) sucrose (grown for 5 extra days), treated with or without 1 μM eBL for 1 hour, were used to total RNA extraction. The eBL (PhytoTechnology Laboratories) was added from a 5 mM stock solution freshly prepared in 80% (v/v) ethanol. Plant tissue was grounded to a fine powder in liquid nitrogen. Approximated 100 mg of ground tissue per sample were homogenized in 1 mL of the commercial reagent TRIsure (Bioline), and total RNA was extracted following the manufacturer’s instructions. The RNA concentration and purity was determined spectrophotometrically (Nanodrop ND-1000 Spectrophotometer). RNA samples (10μg per sample) were DNase-treated with Turbo DNA-free DNase (Ambion) and 1 μg of RNA per sample was run on a 1% agarose gel to confirm RNA integrity. First-strand cDNA was synthesized from 1 μg of RNA by using the iScript cDNA synthesis kit (BioRad), according to the manufacturer’s instructions. cDNAs were amplified in triplicate by quantitative PCR by using SsoFast EvaGreen supermix (BioRad) and the MyiQ Thermal cycler (Bio Rad). The relative expression values were determined by using *ACTINE 2* as a reference gene and plotted relative to Col-0 mock treated expression level. Primers used for quantitative RT-PCR are listed in **Table S2**.

### Transient Expression in *N. benthamiana*

For transient expression in *Nicotiana benthamiana, Agrobacterium tumefaciens* (GV3101::pMP90) carrying the different constructs were used together with the p19 strain ^79^ for infiltration into 4- to 5-week-old *N. benthamiana* leaves at the abaxial side of the leaf lamina. After infiltration, all plants were kept in the greenhouse and analyzed 2 days later. Agrobacteria cultures were grown overnight in LB medium containing rifampicin (50 μg/mL), gentamycin (25 μg/mL) and the construct specific antibiotic. Cells were then harvested by centrifugation (15 minutes, 3000g in 50 mL falcon tubes), pellets were resuspended in agroinfiltration solution (10 mM morpholineethanesulfonic acid (MES) pH 5.6, 10 mM MgCl_2_, and 1 mM acetosyringone) and incubated 2 hours in dark conditions at room temperature. For double infiltration experiments, *Agrobacterium* strains were infiltrated at optical density at 600 (OD_600_) of 0.4 for the constructs and 0.2 for the p19 strain. For triple infiltration experiments, *Agrobacterium* strains were infiltrated at OD_600_ of 0.26 for the constructs and at OD_600_ of 0.2 for the p19 strain. An *Agrobacterium* strain harboring an empty vector (or GUS-HA expressing vector) was used as a negative control to equal the final optical density, in order to obtain a total OD_600_ of approximated 1 in all the infiltration experiments.

For eBL treatment analysis, leaves were pre-treated with 5 μM BL for 3 hours prior to samples collection. *N. benthamiana* leaves were infiltrated with water or 5 μM eBL (PhytoTechnology Laboratories) infiltration solution (10 μL eBL 5mM stock solution in 10 mL H2O), made from a 5 mM stock solutions freshly prepared in 80% (v/v) ethanol.

For brassinazole (BRZ) treatment experiments, the agroinfiltration solution was supplemented with either mock (BRZ solvent as control) or 5 μM BRZ (TCI Europe). After infiltration, *N. benthamiana* plants were kept in the greenhouse and analyzed 2 days later.

### Transient Expression in Arabidopsis *NahG* plants

*Agrobacterium tumefaciens-mediated* expression in Arabidopsis NahG plants ^53^ was performed as described for transient expression in *N. benthamiana* with some modifications. *Agrobacterium* strains were resuspended with an equal OD_600_ in infiltration solution to obtain a total OD_600_ of 0.05 for injection into abaxial leaves side of 4 to 5-week-old Arabidopsis plants. At least 6 plants per co-infiltration mixture and 4 leaves per plant were used per experiment.

### Recombinant Protein Purification and In Vitro Pull-down Assay

The coding sequences of wild-type BRI1 cytoplasmatic domain (residues 814–1196), BRI1 cytoplasmatic domain JMCT9D (residues 250-662), TTL3ΔN1 (residuos 204-691) and TTL3ΔN3 (residuos 567-691) were cloned as described in **Plasmid Constructs** section to generate MBP-BRI1cyt, MBP-BRI1cyt^JMCT9D^, GST-TTL3ΔN1 and GST-TTL3ΔN3 constructs. Recombinant proteins were expressed in *E. coli* strain BL21 (DE3) and extracted using Buffer A (140mM NaCl, 2.7mM KCl, 10mM Na2HPO4, 1.8mM KH2PO4, 1% Triton X-100, pH 8, supplemented with 1 mM PMSF, 0.2 μL/10 mL of Benzonase Nuclease (Sigma), and 1 mg/mL Lisozyme). MBP and GST fusion proteins were purified with Glutathione Sepharose 4B GST-tagged protein purification resin (GE Healthcare) or MBP binding protein coupled to agarose beads (MBP-Trap_A, Chromotek), respectively, according to the manufactures.

To investigate protein-protein interactions, the GST-tagged proteins were first capture by the glutathione agarose-coated beads and then incubated with the MBP-tagged proteins in dilution/wash buffer [50 mM Tris-HCl, pH 7.5; 150 mM NaCl; 10% glycerol; 10 mM EDTA, pH 8; 10 mM DTT; 0,5 mM PMSF; 1% (v/v) P9599 protease inhibitor cocktail (Sigma)] at 4°C during 1 hour in a end-over-end rocker. Protein-protein interaction complexes bound to the glutathione agarose-coated beads were pulled down, washed three times with the dilution/wash buffer and analyzed by western blot as described in the **western blot** section.

Immunoblotted GST and MBP-tagged protein were detected using an anti-GST antibody (Sigma G7781; Dilution 1:10000) and a specific anti-BRI1 antibody ^71^ (Dilution 1:2000) as described in the “**Western Blot Analysis**” section.

### Protein extraction and Co-Immunoprecipitation in *N. benthamiana*

Protein extraction and Co-Immunoprecipitation in *N. benthamiana* were performed as previsouly described ^80^ with some modifications. Briefly, Four-week-old *N. benthamiana* plants were used for transient expression assays as described in **Transient expression in *N. benthamiana*** section. Leaves were grounded to fine powder in liquid nitrogen. Approximated 0,5g of grounded leaves per sample were used and total proteins were then extracted with extraction buffer [50 mM Tris-HCl, pH 7.5; 150 mM NaCl; 10% glycerol; 10 mM EDTA, pH 8; 1 mM NaF; 1 mM Na_2_MoO_4_·2H_2_O; 10 mM DTT; 0,5 mM PMSF; 1% (v/v) P9599 protease inhibitor cocktail (Sigma); Nonidet P-40, CAS: 903619-5 (USB Amersham life science) 0,5% (v/v) for CoIP involving transmembrane proteins BRI1 and BAK1, and 0,2% (v/v) for the rest of CoIP] added at 2 mL/g of powder using an end-over-end rocker during 30 minutes at 4°C. Samples were centrifuged 20 minutes at 4 °C and 9000 g. Supernatants (approximated 4 mg/mL protein) were filtered by gravity through Poly-Prep Chromatography Columns (#731-1550 Bio-Rad) and 100 μL were saved to analyze by western blot as input. The remaining supernatants were incubated 2 hours at 4 °C with 15 μL GFP-Trap coupled to agarose beads (Chromotek) in an end-over-end rocker. During incubation of protein samples with GFP-Trap beads the final concentration of detergent (Nonidet P-40) was adjusted to 0,2% (v/v) in all cases to avoid unspecific binding to the matrix as recommended by the manufacturer. Following incubation, the beads were collected and washed four times with the wash buffer (similar to extraction buffer but without detergent). Finally, beads were resuspended in 75 μL of 2x concentrated Laemmli Sample Buffer and heated at 60°C 30 minutes (for CoIP involving transmembrane proteins BRI1 and BAK1) or (70°C for 20 minutes (for the rest of CoIPs) to dissociate immunocomplexes from the beads. Total (input), immunoprecipitated (IP) and Co-Immunoprecipitated (CoIP) proteins were separated in a 10% SDS-PAGE gel, and analyzed as described in the **Western Blot Analysis** section.

### Bimolecular Fluorescence Complementation (BiFC) Assays

Leaves were co-agroinfiltrated as described in the **Agrobacterium-Mediated Transient Expression in *Nicotiana benthamiana*** section with the *Agrobacterium* strain harboring a construct to express a given protein (Protein A) fused to the N-terminus half of the YFP (Protein A-nYFP) and the BiFC partner protein (Protein B) fused to the C-terminus half of the YFP (Protein B-cYFP), and the other way around (Protein A-cYFP and Protein B-nYFP) to test both BiFC directions. Leaves were observed under the confocal microscope two days after infiltration, as described in **Confocal Imaging of Arabidopsis and *Nicotiana benthamiana*** section.

### Confocal Imaging of Arabidopsis and *N. benthamiana*

Arabidopsis seedlings were germinated in half-strength Murashige-Skoog agar solidified medium (1 % agar (w/v) for vertical growth) supplemented with 1.5% sucrose. For eBL treatment analysis, 4-day-old seedling were incubated in 2 mL of half-strength Murashige and Skoog medium supplemented with 1,5% (w/v) sucrose containing either mock (eBL solvent as control) or 1 μM eBL (PhytoTechnology Laboratories). For BRZ/eBL treatment analysis, seedling with three days and an half were incubated in 2 mL of half-strength Murashige and Skoog medium supplemented with 1% (w/v) sucrose, containing either mock (BRZ solvent as control) or 5 μM BRZ (TCI Europe) for 12 hours (overnight).

The next morning samples were further treated with mock or 1 μM eBL (PhytoTechnology Laboratories) for another 1 hour before being analyzed by confocal microcopy. The eBL (PhytoTechnology Laboratories) and BRZ (TCI Europe) were added from a 5 mM stock solutions freshly prepared in 80% (v/v) ethanol. For visualizations of plasma membrane, seedlings were incubated in 1 μL ddH_2_O containing 1 μg/mL FM4-64 (Invitrogen Molecular Probes) prepared from a 1 mg/mL stock solution for 3-4 minutes, rinsed in ddH_2_O to remove the excess of stain and visualized under confocal microscopy.

For confocal imaging of *Nicotiana benthamiana* leaves in co-expression and BiFC experiments, GFP or YFP fluorescence of the lower epidermis of leaf was visualized with the confocal 2 days after infiltration.

Confocal imaging of Arabidopsis *NahG* plants was performed as described for *Nicotiana benthamiana*, but in this case, images are a maximum Z-projection of seven 1 μm spaced confocal planes from the cell equatorial plane to the cell surface.

All confocal images were obtained using a Leica TCS SP5 II confocal microscope equipped with a 488-nm argon laser for GFP and YFP, and a 561nm He-Ne laser for FM4-64. Leica LAS AF Lite platform and the Java-based image-processing program FIJI ^81,82^ were used in the processing of all microscopy images.

### Stereo Microscopy of Arabidopsis Seedlings

Representative images of Arabidopsis seedlings were acquired using the Nikon Eclipse Ti basic Fluorescence Microscope system with filter for GFP. Wilde-type Col-0 Arabidopsis seedlings were used as negative control for GFP autoflorescence.

### GUS Staining Assay

Four-day-old seedlings were transferred to a medium containing 0,2 μM eBL (PhytoTechnology Laboratories) during 24 hours and then stained for GUS activity. Plant tissues were immersed in histochemical GUS staining buffer (100mM NaPO_4_ pH7, 0.5 mM Ks[Fe(CN)a], 0.5 mM K_4_[Fe(CN)e], 20% Methanol, 0.3% Triton X-100 and 2 mM 5-Bromo-4-chloro-3-indoxyl-beta-D-glucuronide cyclohexylammonium (X-gluc) (Gold Biotechnology, USA)) on multi-well plates, vacuum-infiltrated (60 cm Hg) for 10 minutes three times, and then wrapped in aluminum foil and incubated at 37°C for 12 hours. Samples were then washed several times with 95% ethanol until complete tissue clarification, stored in 50% glycerol and photographed using the Nikon AZ100 Multizoom microscope system.

### Protoplasts Transient Expression Assays

Protoplasts extraction and transfection was performed as previously described ^83^ Briefly, leaves from 5-week-old Arabidopsis Col-0 grown at 10-hour daylight photoperiod were cut to strips and digested for 3 hours in the darkness at room temperature. Protoplasts were then washed and resuspended to a concentration of 5×10^5^ protoplasts/mL before PEG-mediated transfection for 10 minutes. Twenty microliters μl of plasmid expressing GFP or 100 μl of plasmids expressing TTL3-GFP/BZR1-HA were used to transfect 2 mL protoplasts for each transfection. All the plasmids were used at a concentration of 1 μg/ul. The transfected protoplasts were incubated for 6 hours at room temperature and collected for protein extraction and immunoprecipitation, as described for *N. benthamiana* samples.

### Yeast Two-Hybrid Assay

The Gal4-based yeast two-hybrid system (Clontech Laboratories Inc.) was used for testing the interaction between TTL3 and different components of the brassinosteroid signalling pathway. The bait and prey constructs are explained in the “**Plasmid Constructs**” section. The bait and prey plasmids were transformed into *Saccharomyces cerevisiae* strain AH109 as previously described ^84^ and transformants were grown on plasmid-selective media (SD/-Trp-Leu). Plates were incubated at 28 °C for 4 days and independent colonies for each bait-prey combination were resuspended in 200 μl of sterile water. 10-fold serial dilutions were made and 5 μl of each dilution were spotted onto three alternative interaction-selective medium (SD/-Trp-Leu-His+3-AT (3-amino-1, 2, 4-triazole, 2mM), SD/-Trp-Leu-Ade, and SD/-Trp-Leu-Ade+3-AT). Plates were incubated at 28 °C and photographed 3 or 7 days later.

### Yeast Two-Hybrid Protein Extraction

For inmunoblot analysis, one or two independent yeast co-transformants (a and b) for each bait-prey plasmid combination were grown in 50 mL of SD/-Leu-Trp to an OD600 of 0.7-1. Cultures were centrifuged at 4.000 rpm for 3 minutes.

The resulting pellet was washed once with cold water and resuspended in 200 μl of RIPA buffer (2 mM sodium phosphate buffer pH 7, 0,2% Triton X-100, 0,02%-w/v-SDS, 0,2 mM EDTA pH 8, 10 mM ClNa) containing protease inhibitor (1 tablet/10mL, cOmplete, Mini, EDTA-free Protease Inhibitor Cocktail, Roche) Glass beads (500 μl, 425-600 um, Sigma) were added and the sample was vortexed in FastPrepTM FP120 (BIO 101) at power setting 5.5 for two 15 seconds intervals separated by 1 minute intervals on ice. Then 400 μl RIPA buffer with protease inhibitors were added and the sample was vigorously vortexed. The supernatant was recovered, and the protein concentration was determined using Bradford assays. Total protein (50 μg) was resolved on 10% polyacrylamide/SDS gels and analyzed by immunoblotting using a anti-Myc Tag (1:2000, Abgent) which is transcriptionally fused to Gal4BD, as described in the “**Western Blot Analysis**” section

### Extraction of Total Protein from Arabidopsis

Arabidopsis tissue was grounded to fine powder in liquid nitrogen. Approximated 100 mg of grounded tissue per sample were used for total proteins extraction. Denatured protein extracts were obtained by homogenizing and incubating plant material in 200 μL of 2X Laemmli buffer [125 mM Tris-HCl pH 6.8; 4% (w/v) SDS; 20% (v/v) Glycerol; 2% (v/v) Beta-mercaptoethanol; 0, 01% (w/v); Bromophenol blue] for 5 minutes at 95°C, centrifuged (5 minutes, 20 000 g) and the total proteins from supernatant were separated in a 10% SDS-PAGE gel, and analyzed as described in the **Western Blot Analysis** section.

### Western Blot Analysis

Proteins separated by SDS-PAGE polyacrylamide gel electrophoresis were electroblotted using Trans-blot Turbo Transfer System (BioRAD) onto polyvinylidene difluoride (PVDF) membranes (Immobilon-P; Millipore) following instructions by the manufacturer (preprogramed protocols optimized for the molecular weight of the proteins of interest). PVDF membranes, containing electroblotted proteins, were then incubated with the appropriate primary antibody followed by the appropriate secondary second peroxidase-conjugated antibody. In addition to the primary antibodies described in the previous methods section, the following primary antibodies were used for detection of epitope-tagged proteins: mouse monoclonal anti-GFP clone B-2 (1:600; Santa Cruz Biotechnology); mouse monoclonal anti-HA clone HA-7 (1:3000; Sigma-Aldrich); rabbit polyclonal anti-mCherry (1:3000 GeneTex). The secondary antibodies used in the present study were: anti-mouse IgG whole molecule-Peroxidase (1:80000; Sigma-Aldrich) and anti-rabbit IgG whole molecule-Peroxidase (1:14000 or 1:80000; Sigma-Aldrich)

Proteins and epitope-tagged proteins on immunoblots were detected by using the Clarity ECL Western Blotting Substrate or SuperSignal West Femto Maximum Sensitivity Substrate according to the manufacturer’s instructions, and images of different time exposures were acquired by using the Chemidoc XRS+System (Biorad). SDS-PAGE polyacrylamide gels and immunoblotted PVDF membranes were stained with Coomassie blue for confirming equal loading of the different samples in a given experiment.

## QUANTIFICATION AND STATISTICAL ANALYSIS

### Arabidopsis eFP Browser Data Analysis

Gene expression level data from hormone responses was retrieved from Arabidopsis eFP Browser (Hormone Series) web site available from the following link: http://bar.utoronto.ca/efp/cgi-bin/efpWeb.cgi ^85^ Data used for the analysis was obtained from 7-day-old wild-type seedlings. Differential expression was calculated by dividing the expression value of each gene in a given hormone treatment by the corresponding mock control (fold-change of hormone treatment relative to the mock). Hormone gene expression response calculation and Heatmap was obtained using Microsoft Office Excel (Microsoft). Heatmap red colors represent induction and blue colors represent repression as response to the indicated hormone.

### Quantification of Fluorescent Protein Signal

For quantification of fluorescent protein signal in plasma membrane vs cytoplasm all images were analyzed using FIJI software ^81,82^. To measure the ratio between nuclear and cytoplasmic signals, a small area of fixed size (8 pixels) was drawn, and measurements of integrated densities were taken from representative areas within the plasma membrane and cytoplasm of each cell. To delimitate de plasma membrane area, FM4-64 was used to stain the cells. Average ratios between plasma membrane and cytoplasmic signal intensities were calculated based on measures from 3 cells per plant. n=10 plants analyzed (3 cells per plant). This experiment was repeated twice with similar results.

Additionally, for quantification of fluorescent protein signal, lines scan measurements spanning membrane and cytoplasm were carried out from images using FIJI ^81,82^ software, and representative plot profiles of sample measurements are presented in **Figure 3I**.

### Statistics

Band intensity quantification of protein signal detected by western blot, integrated densities from representative areas within the plasma membrane and cytoplasm of each cell analyzed by confocal imaging, as well as Arabidopsis root and hypocotyl lengths were measured from images using FIJI ^81,82^ software. The data for qRT-PCR were gathered with MyiQ optical system software (Bio Rad). For statistical analysis unpaired t-test was performed using GraphPad Prism version 6.00 for Mac (GraphPad Software, La Jolla California USA, www.graphpad.com). Asterisks indicate statistical differences between mutant vs Col-0, unless otherwise specified, as determined by the unpaired *t-test* (* P ≤ 0.05, ** P ≤ 0.01, *** P ≤ 0.001 **** P ≤ 0.0001). Data represent mean values, error bars are SEM. In figure legends, n means number of plants for phenotypic analysis, numbers of biological replicates (3 technical replicates per biological replicate) for qRT-PCR analysis, or number of cells (3 independent measurements per performed per cell) analyzed for quantification of fluorescent protein signal in plasma membrane vs cytoplasm. The experiments were repeated at least three times with similar results.

### *In silico* Three-Dimensional Structural Model of TTL3

The *in silico* protein structure prediction for TTL3 protein was built by submitting primary sequences to the I-TASSER server ^69^ and processed by PyMOL (Schrödinger). Intrinsically disordered regions (IDRs) were predicted using GlobPlot 2, available in the web page (http://globplot.embl.de/).

Tetratricopeptide Repeat (TPR) and thioredoxin-like (TPRX) domains were predicted using SMART/Pfam server and were previously described ^12^.

## Acknowledgements

This work was supported the Ministerio de Economía y Competitividad (cofinanced by the European Regional Development Fund; grants no. BIO2014-55380-R and BIO2017-82609-R to M.A.B) and by a Formación del Personal Investigador Fellowship from the Ministerio de Economía y Competitividad (FPI-BES 2015-071256 to A. G-M). A.P.M. was funded by the Shanghai Center for Plant Stress Biology (Chinese Academy of Sciences) and the Chinese 1000 Talents Program, and by grants from the Gatsby Charitable Foundation and the European Research Council (grant ‘PHOSPHinnATE’) to C.Z., while working in the C.Z. laboratory.

We thank Yanhai Yin and Michael Hothorn for generously providing the anti-BES1 and the anti-BRI1 antibodies, respectively.

We are grateful to Salomé Prat (*BIN2* and *BES1* expression clones), Santiago Mora Garcia (BSU expression clone), José Alonso (GFP-tagged GW vector for expression in protoplasts) and Xiaofeng Wang (pMAC-flag-BRI1-CD-JMCT9D, BRI1 phosphomimetic mutant) for providing expression clones and vectors used in the present study.

## Author contributions

All authors designed the experiments. V.A-S., A.G-M., A.C., N.L., J.P-S., Y.L., A.E-V., D.P., J.P-R., and A.P.M. performed the experiments and analyzed the data. V.A-S., A.P.M., and M.A.B. wrote the manuscript. All authors commented on the manuscript.

## REFERENCES

1. Meldau, S., Erb, M. & Baldwin, I. T. Defence on demand: mechanisms behind optimal defence patterns. Ann. Bot. 110, 1503–1514 (2012).

2. Chaiwanon, J., Wang, W., Zhu, J.-Y., Oh, E. & Wang, Z.-Y. Information Integration and Communication in Plant Growth Regulation. Cell 164, 1257–1268 (2016).

3. Belkhadir, Y., Yang, L., Hetzel, J., Dangl, J. L. & Chory, J. The growth-defense pivot: crisismanagement in plants mediated byLRR-RK surface receptors. Trends Biochem Sci 39, 447–456 (2014).

4. Jaillais, Y. & Vert, G. Brassinosteroid signaling and BRI1 dynamics went underground. Curr. Opin. Plant Biol. 33, 92–100 (2016).

5. Belkhadir, Y. & Jaillais, Y. The molecular circuitry of brassinosteroid signaling. New Phytol. 206, 522–540 (2015).

6. Lozano-Durán, R. & Zipfel, C. Trade-off between growth and immunity: role of brassinosteroids. Trends Plant Sci. 20, 12–19 (2015).

7. Nolan, T. M. et al. Selective Autophagy of BES1 Mediated by DSK2 Balances Plant Growth and Survival. Dev. Cell 41, 33–46.e7 (2017).

8. Zhang, Z. et al. TOR Signaling Promotes Accumulation of BZR1 to Balance Growth with Carbon Availability in Arabidopsis. Curr. Biol. 26, 1854–1860 (2016).

9. Tian, Y. et al. Hydrogen peroxide positively regulates brassinosteroid signaling through oxidation of the BRASSINAZOLE-RESISTANT1 transcription factor. Nat Commun 9, 1063–13 (2018).

10. Wang, W., Bai, M.-Y. & Wang, Z.-Y. The brassinosteroid signaling network-a paradigm of signal integration. Curr. Opin. Plant Biol. 21, 147–153 (2014).

11. Rosado, A. et al. The Arabidopsis tetratricopeptide repeat-containing protein TTL1 is required for osmotic stress responses and abscisic acid sensitivity. Plant Physiol. 142, 1113–1126 (2006).

12. Lakhssassi, N. et al. The Arabidopsis tetratricopeptide thioredoxin-like gene family is required for osmotic stress tolerance and male sporogenesis. Plant Physiol. 158, 1252–1266 (2012).

13. Ceserani, T., Trofka, A., Gandotra, N. & Nelson, T. VH1/BRL2 receptor-like kinase interacts with vascular-specific adaptor proteins VIT and VIK to influence leaf venation. The Plant Journal 57, 1000–1014 (2009).

14. Prasad, B. D., Goel, S. & Krishna, P. In silico identification of carboxylate clamp type tetratricopeptide repeat proteins in Arabidopsis and rice as putative co-chaperones of Hsp90/Hsp70. PLoS ONE 5, e12761 (2010).

15. Samakovli, D., Margaritopoulou, T., Prassinos, C., Milioni, D. & Hatzopoulos, P. Brassinosteroid nuclear signaling recruits HSP90 activity. New Phytol. 203, 743–757 (2014).

16. Shigeta, T. et al. Heat shock protein 90 acts in brassinosteroid signaling through interaction with BES1/BZR1 transcription factor. J. Plant Physiol. 178, 69–73 (2015).

17. Shigeta, T. et al. Molecular evidence of the involvement of heat shock protein 90 in brassinosteroid signaling in Arabidopsis T87 cultured cells. Plant Cell Rep. 33, 499–510 (2014).

18. Lachowiec, J. et al. The protein chaperone HSP90 can facilitate the divergence of gene duplicates. Genetics 193, 1269–1277 (2013).

19. Yang, C.-J., Zhang, C., Lu, Y.-N., Jin, J.-Q. & Wang, X.-L. The mechanisms of brassinosteroids’ action: from signal transduction to plant development. Mol Plant 4, 588–600 (2011).

20. Blatch, G. L. & Lässle, M. The tetratricopeptide repeat: a structural motif mediating protein-protein interactions - Blatch - 1999 - BioEssays - Wiley Online Library. Bioessays (1999).

21. D’Andrea, L. D. & Regan, L. TPR proteins: the versatile helix. Trends Biochem Sci 28, 655–662 (2003).

22. Yang, J. et al. Molecular basis for TPR domain-mediated regulation of protein phosphatase 5. EMBO J 24, 1–10 (2005).

23. Caño-Delgado, A. et al. BRL1 and BRL3 are novel brassinosteroid receptors that function in vascular differentiation in Arabidopsis. Development 131, 5341–5351 (2004).

24. Habchi, J., Tompa, P., Longhi, S. & Uversky, V. N. Introducing Protein Intrinsic Disorder. Chem. Rev. 114, 6561–6588 (2014).

25. Ma, X., Xu, G., He, P. & Shan, L. SERKing Coreceptors for Receptors. Trends Plant Sci. 21, 1017–1033 (2016).

26. Gou, X. et al. Genetic evidence for an indispensable role of somatic embryogenesis receptor kinases in brassinosteroid signaling. PLoS Genet 8, e1002452 (2012).

27. He, K. et al. BAK1 and BKK1 regulate brassinosteroid-dependent growth and brassinosteroid-independent cell-death pathways. CURBIO 17, 1109–1115 (2007).

28. Chinchilla, D. et al. A flagellin-induced complex of the receptor FLS2 and BAK1 initiates plant defences. Nature 448, 497–500 (2007).

29. Schwessinger, B. et al. Phosphorylation-Dependent Differential Regulation of Plant Growth, Cell Death, and Innate Immunity by the Regulatory Receptor-Like Kinase BAK1. PLoS Genet 7, e1002046 (2011).

30. van Esse, W., van Mourik, S., Albrecht, C., van Leeuwen, J. & de Vries, S. A Mathematical Model for the Coreceptors SOMATIC EMBRYOGENESIS RECEPTOR-LIKE KINASE1 and SOMATIC EMBRYOGENESIS RECEPTOR-LIKE KINASE3 in BRASSINOSTEROID INSENSITIVE1-Mediated Signaling. Plant Physiol. 163, 1472–1481 (2013).

31. Du, J. et al. Somatic Embryogenesis Receptor Kinases Control Root Development Mainly via Brassinosteroid-Independent Actions in Arabidopsis thaliana. J Integrative Plant Biology 54, 388–399 (2012).

32. Bernardo-García, S. et al. BR-dependent phosphorylation modulates PIF4 transcriptional activity and shapes diurnal hypocotyl growth. Genes Dev. 28, 1681–1694 (2014).

33. Zhang, Y. et al. Brassinosteroid is required for sugar promotion of hypocotyl elongation in Arabidopsis in darkness. Planta 242, 881–893 (2015).

34. Li, J. et al. BAK1, an Arabidopsis LRR receptor-like protein kinase, interacts with BRI1 and modulates brassinosteroid signaling. Cell 110, 213–222 (2002).

35. Nam, K. H. & Li, J. BRI1/BAK1, a receptor kinase pair mediating brassinosteroid signaling. Cell 110, 203–212 (2002).

36. Wang, R. et al. The Brassinosteroid-Activated BRI1 Receptor Kinase Is Switched off by Dephosphorylation Mediated by Cytoplasm-Localized PP2A B’ Subunits. Mol Plant 9, 148–157 (2016).

37. Lin, W. et al. Inverse modulation of plant immune and brassinosteroid signaling pathways by the receptor-like cytoplasmic kinase BIK1. Proc. Natl. Acad. Sci. U.S.A. 110, 12114–12119 (2013).

38. Tanaka, K. et al. Brassinosteroid homeostasis in Arabidopsis is ensured by feedback expressions of multiple genes involved in its metabolism. Plant Physiol. 138, 1117–1125 (2005).

39. Vriet, C., Russinova, E. & Reuzeau, C. From squalene to brassinolide: the steroid metabolic and signaling pathways across the plant kingdom. Mol Plant 6, 1738–1757 (2013).

40. Chung, Y. & Choe, S. The Regulation of Brassinosteroid Biosynthesis in Arabidopsis. Critical Reviews in Plant Sciences 32, 396–410 (2013).

41. Wilma van Esse, G. et al. Quantification of the brassinosteroid insensitive1 receptor in planta. Plant Physiol. 156, 1691–1700 (2011).

42. Fàbregas, N. et al. The brassinosteroid insensitive1-like3 signalosome complex regulates Arabidopsis root development. Plant Cell 25, 3377–3388 (2013).

43. Geldner, N., Hyman, D. L., Wang, X., Schumacher, K. & Chory, J. Endosomal signaling of plant steroid receptor kinase BRI1. Genes Dev. 21, 1598–1602 (2007).

44. Vida, T. A. A new vital stain for visualizing vacuolar membrane dynamics and endocytosis in yeast. The Journal of Cell Biology 128, 779–792 (1995).

45. Wang, X. & Chory, J. Brassinosteroids regulate dissociation of BKI1, a negative regulator of BRI1 signaling, from the plasma membrane. Science 313, 1118–1122 (2006).

46. Maselli, G. A. et al. Revisiting the evolutionary history and roles of protein phosphatases with Kelch-like domains in plants. Plant Physiol. 164, 1527–1541 (2014).

47. Vert, G. & Chory, J. Downstream nuclear events in brassinosteroid signalling. Nature 441, 96–100 (2006).

48. Gampala, S. S. et al. An essential role for 14-3-3 proteins in brassinosteroid signal transduction in Arabidopsis. Dev. Cell 13, 177–189 (2007).

49. Ryu, H. et al. Nucleocytoplasmic shuttling of BZR1 mediated by phosphorylation is essential in Arabidopsis brassinosteroid signaling. Plant Cell 19, 2749–2762 (2007).

50. Shimada, S. et al. Formation and dissociation of the BSS1 protein complex regulates plant development via brassinosteroid signaling. THE PLANT CELL ONLINE 27, 375–390 (2015).

51. He, J. X., Gendron, J. M., Yang, Y., Li, J. & Wang, Z. Y. The GSK3-like kinase BIN2 phosphorylates and destabilizes BZR1, a positive regulator of the brassinosteroid signaling pathway in Arabidopsis. Proc. Natl. Acad. Sci. U.S.A. 99, 10185–10190 (2002).

52. Kim, T.-W. et al. Brassinosteroid signal transduction from cell-surface receptor kinases to nuclear transcription factors. Nat. Cell Biol. 11, 1254–1260 (2009).

53. Rosas-Diaz, T. et al. Arabidopsis NahG plants as a suitable and efficient system for transient expression using Agrobacterium tumefaciens. Mol Plant (2016). doi:10.1016/j.molp.2016.11.005

54. Sreeramulu, S. et al. BSKs are partially redundant positive regulators of brassinosteroid signaling in Arabidopsis. Plant J 74, 905–919 (2013).

55. Wang, C. et al. Identification of BZR1-interacting proteins as potential components of the brassinosteroid signaling pathway in Arabidopsis through tandem affinity purification. Mol. Cell Proteomics 12, 3653–3665 (2013).

56. González-García, M.-P. et al. Brassinosteroids control meristem size by promoting cell cycle progression in Arabidopsis roots. Development 138, 849–859 (2011).

57. Tang, W. et al. PP2A activates brassinosteroid-responsive gene expression and plant growth by dephosphorylating BZR1. Nat. Cell Biol. 13, 124–131 (2011).

58. Soutourina, J. Transcription regulation by theMediator complex. Nature Publishing Group 1–13 (2017). doi:10.1038/nrm.2017.115

59. Wang, Z.-Y., Bai, M.-Y., Oh, E. & Zhu, J.-Y. Brassinosteroid signaling network and regulation of photomorphogenesis. Annu. Rev. Genet. 46, 701–724 (2012).

60. Yin, Y. et al. BES1 accumulates in the nucleus in response to brassinosteroids to regulate gene expression and promote stem elongation. Cell 109, 181–191 (2002).

61. Li, V. S. W. et al. Wnt Signaling through Inhibition of β-Catenin Degradation inan Intact Axin1 Complex. Cell 149, 1245–1256 (2012).

62. Clevers, H. & Nusse, R. Wnt/b-Catenin Signaling and Disease. Cell 149, 1192–1205 (2012).

63. Mora-Garcia, S. Nuclear protein phosphatases with Kelch-repeat domains modulate the response to brassinosteroids in Arabidopsis. Genes Dev. 18, 448–460 (2004).

64. Peng, P., Yan, Z., Zhu, Y. & Li, J. Regulation of the Arabidopsis GSK3-like kinase BRASSINOSTEROID-INSENSITIVE 2 through proteasome-mediated protein degradation. Mol Plant 1, 338–346 (2008).

65. Kim, T.-W., Michniewicz, M., Bergmann, D. C. & Wang, Z.-Y. Brassinosteroid regulates stomatal development by GSK3-mediated inhibition of a MAPK pathway. Nature 482, 419–422 (2012).

66. Cai, Z. et al. GSK3-like kinases positively modulate abscisic acid signaling through phosphorylating subgroup III SnRK2s in Arabidopsis. Proc. Natl. Acad. Sci. U.S.A. 111, 9651–9656 (2014).

67. Tang, W. et al. BSKs mediate signal transduction from the receptor kinase BRI1 in Arabidopsis. Science 321, 557–560 (2008).

68. Shi, H. et al. BR-SIGNALING KINASE1 Physically Associates with FLAGELLIN SENSING2 and Regulates Plant Innate Immunity in Arabidopsis. THE PLANT CELL ONLINE 25, 1143–1157 (2013).

69. Zhang, Y. I-TASSER server for protein 3D structure prediction. BMC Bioinformatics 9, 40 (2008).

70. Wang, X. et al. Sequential Transphosphorylation of the BRI1/BAK1 Receptor Kinase Complex Impacts Early Events in Brassinosteroid Signaling - ScienceDirect. Dev. Cell (2008).

71. Bojar, D. et al. Crystal structures of the phosphorylated BRI1 kinase domain and implications for brassinosteroid signal initiation. Plant J 78, 31–43 (2014).

72. Yu, X. et al. A brassinosteroid transcriptional network revealed by genome-wide identification of BESI target genes in Arabidopsis thaliana. Plant J 65, 634–646 (2011).

73. McKinney, E. C. & Meagher, R. B. Members of the Arabidopsis actin gene family are widely dispersed in the genome. Genetics 149, 663–675 (1998).

74. Albrecht, C., Russinova, E., Hecht, V., Baaijens, E. & de Vries, S. The Arabidopsis thaliana SOMATIC EMBRYOGENESIS RECEPTOR-LIKE KINASES1 and 2 control male sporogenesis. Plant Cell 17, 3337–3349 (2005).

75. Lozano-Durán, R., Bourdais, G., He, S. Y. & Robatzek, S. The bacterial effector HopM1 suppresses PAMP-triggered oxidative burst and stomatal immunity. New Phytol. 202, 259–269 (2014).

76. Nakagawa, T. et al. Development of series of gateway binary vectors, pGWBs, for realizing efficient construction of fusion genes for plant transformation. J. Biosci. Bioeng. 104, 34–41 (2007).

77. Christian, G., Rainer, W., rg, K. J., Ralf-R, M. & Robert, H. N. New GATEWAY vectors for High Throughput Analyses of Protein-Protein Interactions by Bimolecular Fluorescence Complementation. Mol Plant 2, 1051–1058 (2009).

78. Clough, S. J. & Bent, A. F. Floral dip: a simplified method for Agrobacterium-mediated transformation of Arabidopsis thaliana. Plant J 16, 735–743 (1998).

79. Voinnet, O., Rivas, S., Mestre, P. & Baulcombe, D. An enhanced transient expression system in plants based on suppression of gene silencing by the p19 protein of tomato bushy stunt virus. Plant J 33, 949–956 (2003).

80. Kadota, Y., Macho, A. P. & Zipfel, C. in Methods in Molecular Biology 1363, 133–144 (Springer New York, 2016).

81. Schneider, C. A., Rasband, W. S. & Eliceiri, K. W. NIH Image to ImageJ: 25 years of image analysis. Nat. Methods 9, 671–675 (2012).

82. Schindelin, J. et al. Fiji: an open-source platform for biological-image analysis. Nat. Methods 9, 676–682 (2012).

83. Yoo, S.-D., Cho, Y.-H. & Sheen, J. Arabidopsis mesophyll protoplasts: a versatile cell system for transient gene expression analysis. Nat Protoc 2, 1565–1572 (2007).

84. Gietz, R. D. & Schiestl, R. H. Transforming yeast with DNA.(Invited chapter). Method Mol. Cell. Biol 5, 255–269 (1995).

85. Winter, D. et al. An ‘Electronic Fluorescent Pictograph’ browser for exploring and analyzing large-scale biological data sets. PLoS ONE 2, e718 (2007).

